# Neuroinflammation in neuronopathic Gaucher disease: Role of microglia and NK cells

**DOI:** 10.1101/2022.05.13.491834

**Authors:** Chandra Sekhar Boddupalli, Shiny Nair, Glenn Belinsky, Joseph Gans, Erin Teeple, Tri-Hung Nguyen, Sameet Mehta, Lilu Guo, Martin L Kramer, Jiapeng Ruan, Hongge Wang, Matthew Davison, D.J Vidyadhara, Zhang Bailin, Katherine Klinger, Pramod K. Mistry

**Affiliations:** Department of Internal Medicine, Yale School of Medicine, New Haven, CT; Translational Sciences, Sanofi, Framingham, MA; Yale Center for Genome Analysis; Department of Neuroscience, Yale School of Medicine

**Keywords:** *Gba*, microglial dysfunction, neuroinflammation, neurodegeneration, glucosylphingosine, lipid.

## Abstract

**Background:** Neuronopathic Gaucher Disease (nGD) is a rare neurodegenerative disorder caused by biallelic mutations in *Gba*, and buildup of glycosphingolipids in lysosomes. Neuronal injury and cell death are prominent pathological features, however the role of *Gba* in individual cell types and involvement of microglia, blood derived macrophages and immune infiltrates in nGD pathology remains enigmatic.

**Methods:** Here, using single cell resolution of mouse nGD brains, we found induction of neuroinflammation pathways involving microglia, NK cells, astrocytes, and neurons.

**Results:** Targeted rescue of *Gba* in microglia and in neurons, respectively in *Gba* deficient, nGD mice reversed the buildup of glucosylceramide (GlcCer) and glucosylsphingosine (GlcSph), concomitant with amelioration of neuroinflammation, reduced level of serum neurofilament light chain (Nf-L) and improved survival. The levels of bioactive lipid, GlcSph was strongly correlated with serum Nf-L and ApoE in nGD mouse models as well as GD patients. *Gba* rescue in microglia/macrophage compartment prolonged survival, that was further enhanced upon treatment with brain permeant inhibitor of glucosylceramide synthase, effects mediated via improved glycosphingolipid homeostasis and reversal of neuroinflammation involving activation of microglia, brain macrophages and NK cells.

**Conclusions:** Together, our study delineates individual cellular effects of *Gba* deficiency in nGD brains, highlighting the central role of neuroinflammation driven by microglia activation and the role of brain permeant small molecule glucosylceramide inhibitor in reversing complex multidimensional pathophysiology of nGD. Our findings advance disease biology whilst identifying compelling biomarkers of nGD to improve patient management, enrich clinical trials and illuminate therapeutic targets.

**Funding:** Research grant from Sanofi Genzyme; other support includes R01NS110354.Yale Liver Center P30DK034989, pilot project grant.

## Introduction

In Gaucher disease (GD), biallelic mutations in *GBA1* underlie defective acid β-glucosidase (glucocerebrosidase, GCase) and buildup of the primary substrate, glucosylceramide (GluCer), and its inflammatory metabolite glucosylsphingosine (GlcSph) in the lysosomes (Grabowski et al., 2021). GD is broadly classified into three phenotypic types based on the absence (type 1 GD, GD1) or presence of early onset neurodegeneration (fulminant GD type 2, GD2) and chronic neurodegeneration (GD3). Adults with GD1 have a markedly increased risk of Parkinson’s disease and Lewy body dementia (PD/LBD) (Bultron et al., 2010; Grabowski et al., 2021). Moreover, heterozygous carriers of *GBA1* mutations are at an increased risk of PD/LBD (Aharon-Peretz et al., 2004; Sidransky and Lopez, 2012; Sidransky et al., 2009). Notably, low GCase activity has been found in the brains of sporadic PD patients who do not harbor *GBA* mutations(Gegg et al., 2022). Together, *GBA* variants and GCase deficiency are important determinants of neurodegenerative diseases.

Defective lysosomal acid β-glucosidase in GD leads to the build-up of lysosomal GluCer and GlcSph, lipids which exhibit potent inflammatory and immunogenic activities (Nagata et al., 2017; Nair et al., 2015; Nair et al., 2016; Pandey et al., 2017). Treatment of non-neuronopathic GD1, involves enzyme replacement therapy (ERT) targeting macrophage mannose receptors and substrate reduction therapies (SRT) using inhibitors of glucosylceramide synthase (GCS) (Mistry et al., 2017a; Mistry et al., 2017b; Platt et al., 2018). However, currently, there are no effective therapies for the devastating neurodegenerative sequela of *GBA1* mutations. In GD1, SRT is predicated on the concept that rate of synthesis of GluCer is reduced to match residual enzyme activity due to *GBA* mutations(Platt et al., 2018). However, in neurodegenerative GD2 and GD3, residual glucocerebrosidase activity is profoundly depressed, due to severe *GBA1* mutations. A randomized controlled trial of brain penetrant SRT, N-butyldeoxynojirimycin in patients GD3 was unsuccessful in ameliorating neurological manifestations. However, N-butyldeoxynojirimycin is relatively weak inhibitor of GCS with even more potent inhibitory off-target effects (Schiffmann et al., 2008). Using a more specific and potent GCS inhibitor in chemically induced model of nGD, some disease pathways were ameliorated on bulk RNA analysis, had no effects on inflammatory pathways (Blumenreich et al., 2021).

Microglia are specialized, self-renewing, CNS-resident macrophages that represent the dominant immune cells involved in maintaining CNS homeostasis in cooperation with other CNS cell types such as neurons, astrocytes, and oligodendrocytes. Emerging data from single-cell analysis and GWAS studies of several neurodegenerative diseases have revealed central role of microglia in neurodegeneration (Chen and Colonna, 2021). Although several mechanisms have been proposed to explain *GBA1*-associated neurodegenerative phenotype, most of the studies in GD have focused on the *GBA1* deficit in neurons, while contribution of other brain cell types in driving the disease pathology has been considered minimal. While there has been immunohistochemical evidence that microglia are altered in neuronopathic GD, few studies have focused on the effect of *GBA1* deficiency *per se* on microglia. Further, studies are warranted to investigate the immune landscape of nGD to clearly delineate therapeutic targets Hitherto, studies aimed at understanding the mechanisms of neuroinflammation in neurodegeneration due to *GBA* deficiency have solely relied on bulk cell population analysis, which have hindered the delineation of heterogeneity and complexity of the immune milieu within the brain as well as role of *GBA1* in individual cell types of the brain. Here, we applied an integrated approach by generating novel mice models, lipidomic analyses, scRNA-seq of immune cells, and brain snRNA-seq to delineate temporospatial components of *GBA1*-associated neuroinflammation, including immune cell subsets, activated pathways, as well as probe therapeutic targets and discover novel biomarkers. Importantly, we used both early onset neuronopathic GD mice *Gba*^lnl/lnl^ (loxP-Neo-loxP) mice, with a germ-line deletion of *Gba* (henceforth referred to as nGD mice) which phenocopies human GD2, fulminant neuronopathic GD (Enquist et al., 2007) as well as a new mouse model of microglia-specific *Gba* deletion in *Gba-floxed* mice that mimics the late-onset progression seen commonly in GD patients. To understand the function of Gba in microglial and neuronal homeostasis and disease progression we also developed two additional new nGD mouse models with microglia and neuron specific rescue of *Gba*.

We evaluated brain-penetrant inhibitor of GCS as a therapeutic strategy in both early and late onset nGD models to assess its impact on reducing cellular glycosphingolipids, individual components of neuroinflammation and neurodegeneration while elucidating temporospatial cellular events. Our findings define a previously unreported role of glycosphingolipid-laden microglia and macrophages along with NK cells and astrocytes in *Gba*-associated neurodegeneration while also revealing novel biomarkers and therapeutic targets.

## Results

### Glycosphingolipid-laden microglial activation and immune cell infiltration drive GD associated neurodegeneration

We performed longitudinal temporospatial analysis of brain in nGD mice (*Gba*^lnl/lnl^ mice with germ-line deletion of *Gba,* rescued from lethal skin phenotype using K14 Cre), compared to control mice (*Gba*^lnl/wt^ and *Gba*^wt/wt^) on days 2, 4, 8, 10, and 14. nGD mice looked clinically well during the first week and subsequently developed progressive ataxia, weight loss, and hind limb paralysis, by day 14, as described previously (Enquist et al., 2007). Microglia and macrophage subsets in brain were defined by combinatorial expression of CD11b, CD45, T cell immunoglobulin and mucin domain containing 4 (TIMD4) and the chemokine receptor C-C motif chemokine receptor 2 (CCR2). Brain microglia have self-renewal capacity with minimal monocyte input in contrast to CCR2^+^ macrophages which have limited self-renewal capacity and are constantly replaced by monocytes (Dick et al., 2022). Infiltration of CCR2^+^ MFs defined as CD11b^hi^ CD45^+^CCR2^+^ CD64^+^ TIMD4^-^ population (fig. S1A) were noted in the nGD brain from an early time point day 2, which showed a steady increase until day 14 when the mice reached the humane endpoint (Fig. 1A). Concurrently, there was attrition of homeostatic microglia and incremental infiltration of diverse immune cells into nGD brains, which coincided with mice displaying clear signs of neurodegeneration (Fig.1A and B and fig. S1A). The predominant immune cells in healthy mouse brains were homeostatic microglia, as expected, whereas in nGD brains, the repertoire of immune cells comprised diverse lymphoid (T cells, NK cells, ILC-2, pDC), and myeloid compartment (cDC, monocytes and/or macrophages) (fig. S1B). To investigate the metabolic consequences of *Gba* deletion in microglia and infiltrating immune cells, we performed HPLC/MS/MS on flow-sorted microglia and infiltrating immune cells. Microglia from nGD brains harbored elevated levels of GluCer species (C16, C18, and C20) as well as GlcSph (fig. S1C). Notably, the immune cell infiltrates in the nGD brains were also enriched in glucosylceramides (C16 and C18) (fig. S1C).

**Fig. 1.**
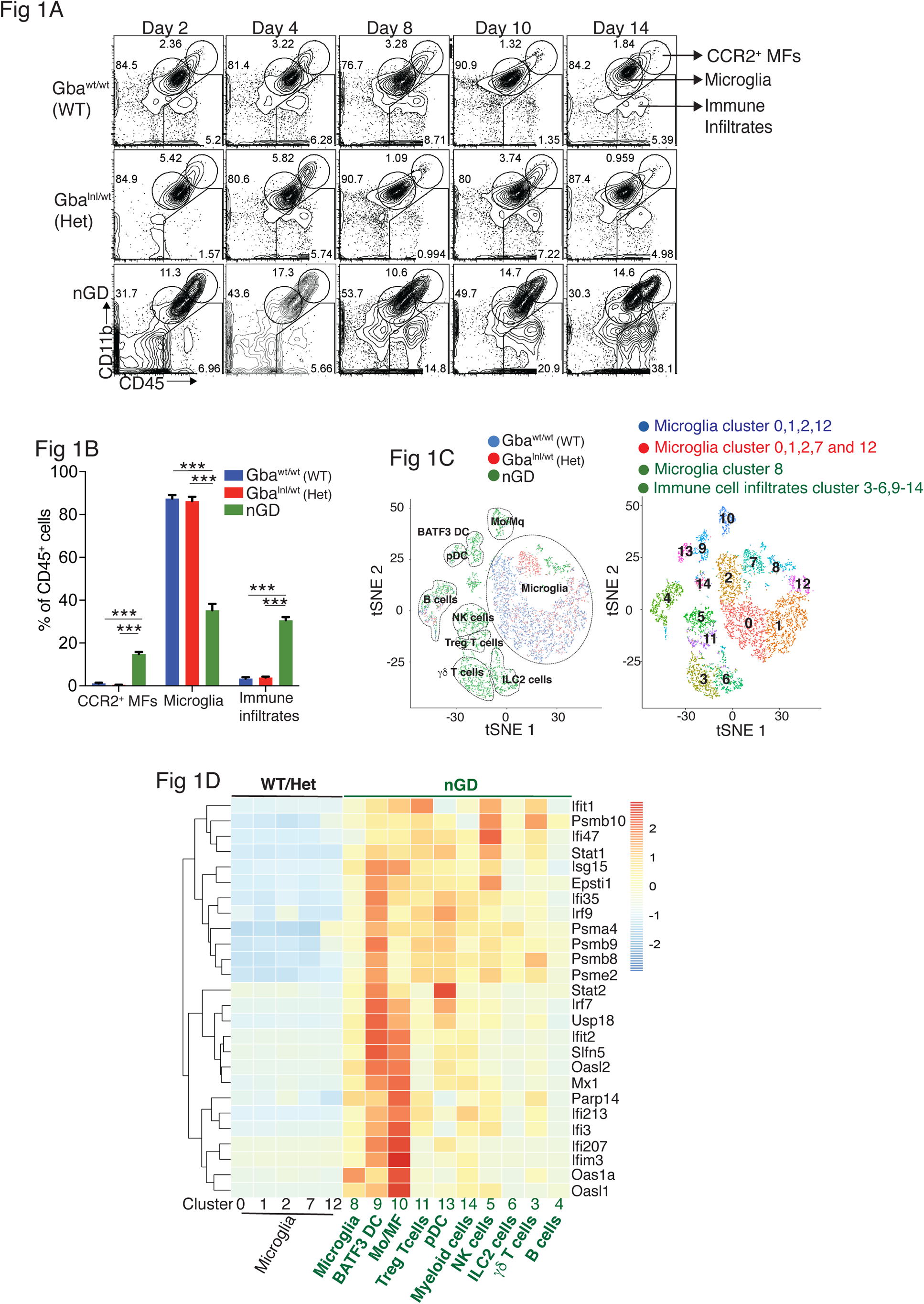
Loss of Gba induces microglial activation and immune cell infiltration in nGD brain. **(A)** FACS analysis on the whole brain of Gba^wt/wt^, Gba^lnl/wt^ and nGD (Gba^lnl/lnl^) mice performed at indicated days. The gates indicate cell populations revealed by CD11b and CD45 expression: CCR2^+^ MFs (CD11b^hi^CD45^+^), microglia (CD11b^lo^CD45^+^) and immune infiltrates (CD11b^lo^CD45^hi^). The data is representative of three independent experiments. (**B)** Bar graph compares percentage of CCR2^+^ MFS, microglia and immune infiltrates between Gba^wt/wt^, Gba^lnl/wt^ and nGD mice brain (n=6 to 8 mice/group); statistical significance was determined using t test with using Bonferroni-Dunn correction for multiple comparisons (***p<0.0001). (**C)** tSNE plot depicting different microglial and non-microglial cell subsets. The clusters are coded based on their mice affiliation (on left). In total, 14 clusters containing 6 microglia clusters and 9 clusters of immune cells (on right). (**D)** Hierarchical clustering of differentially expressed genes associated with type 1 IFN genes from nGD mice versus the control mice. p<0.05 was considered significant (2-sided t tests). All individual type 1 IFN genes with significant differential expression are listed on right. (**H)** Neurofilament light chain (Nf-L) levels in the serum of nGD, nGD Cx3cr1^Cre/+^ and nGD Nes^Cre/+^ mice compared to littermate controls. Experiments are repeated twice, and data presented here is from one experiment (n= 2 to 4 mice/group). Error bars represent Means ± SEM; p values are calculated with Welch’s test (**p<0.001 and ***p<0.0001).

For *de novo* characterization of the brain immune cell microenvironment, sorted CD45^+^ cells from nGD and control brains were analyzed by scRNA-seq and tSNE analysis revealed 15 distinct cellular clusters of CD45^+^ cells (numbered 0-14) (Fig. 1C), which were assigned to individual immune subsets based on the expression of known marker genes (Fig. 1C and fig. S1D and E). There were major alterations in the immune cell composition in nGD brains, which was qualitatively and quantitatively dominated by various lymphocyte populations (T cells, ψοT cells, Treg cells, NK cells, and ILC-2), pDC, and a heterogenous myeloid compartment (cDC, monocytes/macrophages and granulocytes) (Fig. 1C and fig. S1D). Homeostatic microglia were reduced in nGD brains, consistent with impaired microglial homeostasis in nGD pathology (fig. S1A and D). Interestingly, the B cell population (cluster 4) appeared more pronounced in nGD brains, although it was also present in controls. The microglia and immune cell infiltrates in nGD brains exhibited a striking upregulation of type 1-interferon signature genes (ISG) (Fig. 1D). Total RNA-seq analysis performed on flow-sorted microglia from nGD and control mice revealed distinct gene expression profiles in GluCer/GlcSph-laden nGD microglia vs. control microglia (fig. S2A). Microglia of nGD mice exhibited clear downregulation of homeostatic genes and concomitant upregulation of disease-associated microglia (DAM) signature genes, in addition to upregulation of ISGs (fig. S2B and C). Together, these findings establish GluCer/GlcSph-laden DAM, and peripheral immune cell infiltration as key features of nGD neuropathology.

### *Gba* restoration in microglia and neurons prolong survival of nGD mice

To dissect the contribution of microglia and neurons nGD neurodegeneration, we generated nGD Cx3cr1^Cre/+^ and nGD Nes^Cre/+^ mice for selective rescue of *Gba* in microglia and neurons, respectively (Fig. 2A and fig. S3A). Unless otherwise stated, we used littermate *Gba*^wt/wt^ with the respective Cre recombinases as controls. Notably, restoration of *Gba* in microglia (nGD Cx3cr1^Cre/+^ mice) led to more than 2-fold increase in survival compared to nGD mice (Fig. 2B). Selective restoration of *Gba* in neurons (nGD Nes^Cre/+^ mice), further enhanced the survival up to ∼200 days where they reached the humane endpoint (Fig. 2B). To assess the alterations in microglia and macrophage phenotypes associated with targeting of *Gba* in microglia/ neurons, we compared the brains of nGD, nGD Cx3cr1^Cre/+^, and nGD Nes^Cre/+^ mice. Restoration of *Gba* in microglia of nGD mice (nGD Cx3cr1^Cre/+^) resulted in increased homeostatic microglia with reduction in CCR2^+^ MFs and peripheral immune cell infiltration compared to nGD brains (Fig. 2C and fig. S3B). In contrast, restoration of *Gba* in neurons of nGD mice (nGD Nes^Cre/+^) blocked attrition of microglia as well as the infiltration of CCR2+ MFs and peripheral immune cells (Fig. 2C). Correspondingly, induction of intracellular Pro-IL-1ß in microglia, an indicator of microglial activation, was observed in both nGD and nGD Cx3cr1^Cre/+^ brains (Fig 2 D, and E) but not in nGD Nes^Cre/+^ brains (Fig. 2 D and F; fig. S3C). Collectively, these data establish the critical role of *Gba* deficiency in neurons in aiding vigorous microglial activation and immune cell infiltration in nGD and nGD Cx3cr1^Cre/+^ brains. Consistent with this notion, compared to nGD mice, both nGD Cx3cr1^Cre/+^ mice and nGD Nes^Cre/+^ mice also showed significant reduction of brain glucosylceramides (by ≥ 90% for C16 and C18:1, and by 50-70% for C18, C20:1, C22, C22.1, C24, and C24:1, and by 38% for C20 glucosylceramide) with a concomitant, striking reduction of GlcSph (89% and 98%) respectively (Fig. 2G and fig. S4A). As expected, there was no change in the levels of galactosylceramide (GalCer) species or galactosylsphingosine (GalSph) in nGD brains (fig. S4B). MALDI imaging of frozen brain sagittal sections was performed to assess the topography of GSL accumulation. Elevated levels of numerous hexosylceramides (HexCers) were detected in nGD brains. Of these, HexCer (18:1/22:0) exhibited higher signal intensity in the cerebral cortex and midbrain region of nGD Cx3cr1^Cre/+^ brains compared to control mice (fig. S4C). Similarly, HexCer (18:1/20:0) species was elevated in the same brain regions of nGD Nes^Cre/+^ mice (fig. S4D). Interestingly, a co-regulated lipid in GD cells models, lysophosphatidylcholine (LysoPC) (Bodennec et al., 2002), showed striking accumulation in the cerebral cortex and midbrain region of nGD Cx3cr1^Cre/+^ and nGD Nes^Cre/+^ mice (fig. S4C and D lower panel). Assessment of motor balance and coordination showed that the longer-lived adult nGD Nes ^Cre/+^ (4-6 months) mice were unable to complete the balance beam runs with an average time > 60 s (fig. S4E). Notably, there were unusual repetitive circling body movements (fig. S4F). These features appear to phenocopy behavioral disorders reported in human GD3 (Abdelwahab et al., 2017). Taken together, these results indicate the differential role of Gba in individual cell types in the brain. Notably, reconstituting *Gba* in microglia of nGD mice affords an impressive capacity to offset toxic lipid accumulation in the brain and significantly prolong survival.

**Fig. 2.**
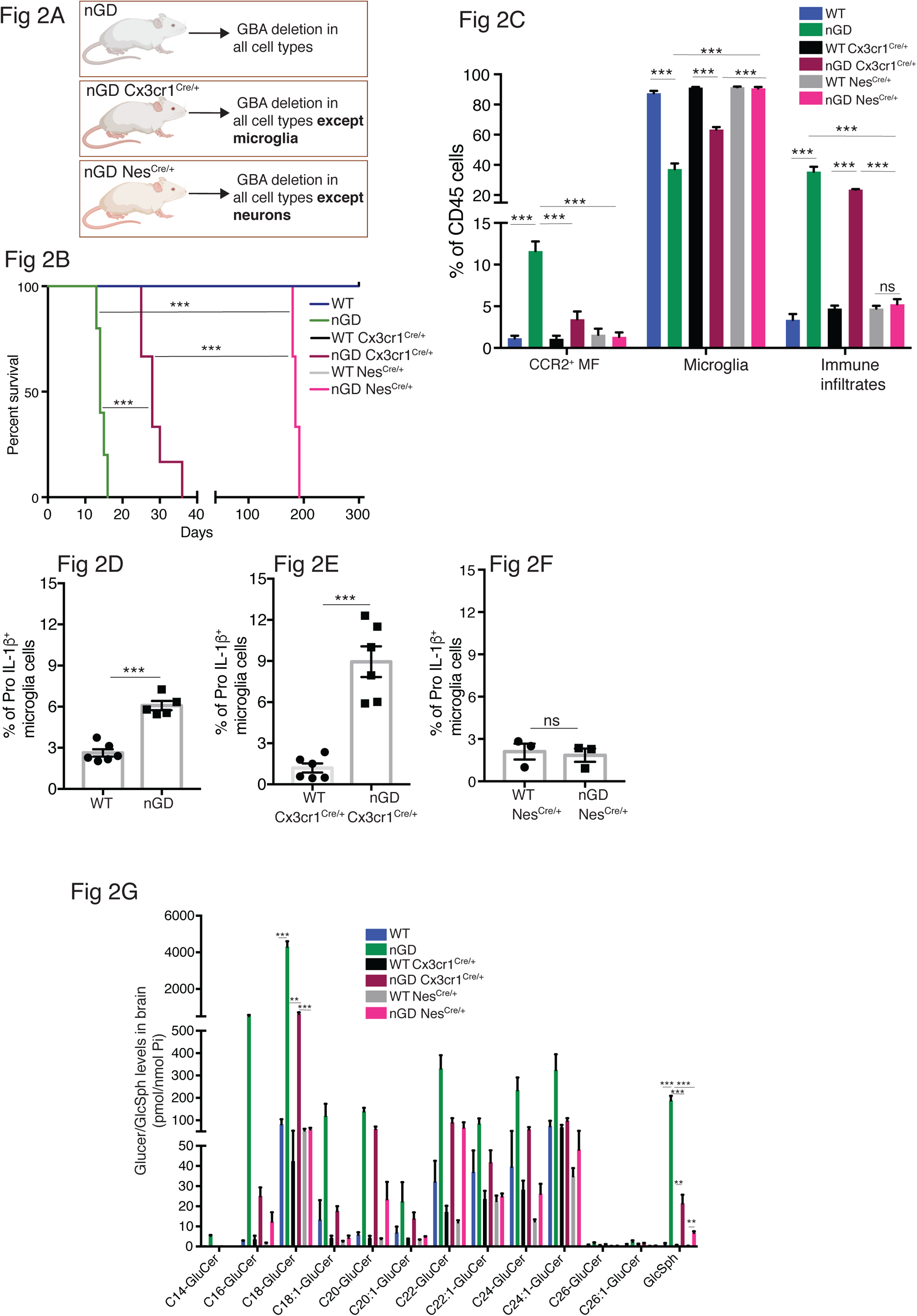
Gba deficiency in neurons aid in microglial activation and immune cell infiltration. **(A)** Schematic showing overview of the mouse models and methods used in the study. (**B)** Kaplan-Meier Survival analysis of nGD, nGD Cx3cr1^Cre/+^ and nGD Nes^Cre/+^ mice cohorts with their respective littermate controls (n= 5 to 6 mice/group) using log-rank (Matel-Cox) test (***p<0.0001). (**C)** Bar graph shows comparison of percentage of CCR2+ MFs, microglia and immune infiltrates between nGD, nGD Cx3cr1^Cre/+^and nGD Nes^Cre/+^ mice with littermate controls (n= 3 to 6 mice/group). Bar graph showing percentage of Pro-IL-1ß^+^ microglia cells in (**D)** nGD vs control mice (n=5-6/ group), (**E)** nGD Cx3cr1^Cre/+^ vs control mice (n=5-6/ group) and (**F)** nGD Nes^Cre/+^ vs the control mice (n=3 mice/group). (**G)** Quantitative analysis of total GluCer species and GlcSph levels by LC-ESI-MS/ MS in nGD, nGD Cx3cr1^Cre/+^ mice and nGD Nes^Cre/+^ mice brain compared with the control mice (n=4-8 mice/group). (**C-G)** shows representative data from two independent experiments using controls. Means ± SEM are shown. Unpaired t-test, two tailed was used to test significance. *p<0.05, **p<0.001 and ***p<0.0001.

### Selective deletion of *Gba* in microglia results late onset neurodegeneration

To further establish the role of *Gba* in microglia, we used our *Gba* floxed mice (Mistry et al., 2010) for microglia-specific *Gba* deletion using Cx3cr1-Cre (Yona et al., 2013) (*Gba*^loxp/loxp^ Cx3cr1^Cre/+^) (fig. S5A). We investigated this mouse strain for clinical phenotype, brain GluCer/GlcSph accumulation, neuroinflammation, and whether microglia/macrophages underwent compensatory recycling in the setting of microglia-specific *Gba* deficiency. Remarkably, *Gba*^loxp/loxp^ Cx3cr1^Cre/+^ brains showed a pronounced accumulation of GlcSph and several GluCer species (C16, C18, C20, and C22 GluCer species) compared to control brains (Fig. 3A). There was no change in the levels of GalCer or GalSph species (fig. S5B). Young mice appeared healthy despite accumulation of GluCer/GlcSph in the brain, but aged *Gba*^loxp/loxp^ Cx3cr1^Cre/+^ mice (∼ 12 months old) started to exhibit motor deficits manifested by a longer time to complete the beam walk and increased tendency to slip (data not shown). HPLC-MS/MS analysis revealed elevated levels of GlcSph in the sera of young *Gba*^loxp/loxp^ Cx3cr1^Cre/+^ mice (6-8 weeks), which tended to increase further in aged (14 months) mice compared to healthy controls (Fig. 3B). Congruently, there was a significant reduction in microglia and increase in CCR2^+^ MFs and immune cell infiltration in aged *Gba*^loxp/loxp^ Cx3cr1^Cre/+^ mice, while no changes were observed in young *Gba*^loxp/loxp^ Cx3cr1^Cre/+^ mice (Fig. 3C). We attributed increase of infiltrating CCR2^+^ MFs in aged *Gba*^loxp/loxp^ Cx3cr1^Cre/+^ mice brain compared to young mice to higher turnover rate as seen by BrdU uptake assay (fig. S5C and D). In contrast to nGD mice with florid neuroinflammation, we could not detect Pro IL-1ß induction in microglia of *Gba*^loxp/loxp^ Cx3cr1^Cre/+^ mice (fig. S5E). We isolated FACS-sorted microglia from young and aged *Gba*^loxp/loxp^ Cx3cr1^Cre/+^ brains to perform scRNAseq. A total of 2,302 microglial cells single-cell transcriptomes were subjected to Louvain clustering, resulting in 9 transcriptionally distinct microglial states (Fig. 3D). The proportion of individual microglial clusters were differentially enriched in young vs aged mice brain (Fig. 3E). Taking advantage of the single-cell resolution of our data and published microglia-specific lineage genes (Chen and Colonna, 2021; Chen et al., 2021; Wang et al., 2020), we identified homeostatic microglial clusters (0, 2, 3, and 6), DAM clusters (1 and 4), a hitherto undescribed *Gba* locus associated gene cluster (cluster 5, we refer to it as *Gba* cluster because *<u>Mtx1</u>* <u>and *Thbs3*</u> are contiguous with *the Gba* gene) and ISG cluster (cluster 7) (Fig. 3E and F). In young *Gba*^loxp/loxp^ Cx3cr1^Cre/+^ mice, homeostatic microglia represented by clusters 0, 2, 3, and 6 were the predominant microglia population (Fig. 3F). Notably, in aged *Gba*^loxp/loxp^ Cx3cr1 ^Cre/+^ brains, DAM signature clusters (1 and 4), along with *the Gba* locus associated gene cluster (cluster 5) and interferon-induced gene cluster (cluster 7) represented the major microglial populations (Fig. 3F). Therefore, with aging transition of homeostatic microglia (cluster 2) into DAM microglia cluster with higher expression of DAM genes (*Apoe*, *Spp1*, *Lpl*, *Ccl3*, and *Cst7*) and lower levels of homeostasis genes (*P2ry12* and *Tmem119*) was evident (Fig. 3E). Aged *Gba*^loxp/loxp^ Cx3cr1^Cre/+^ microglia also expressed higher levels of chemokines (*Ccl3*, *Ccl4*, *Ccl2*, etc.), inflammatory molecules (*Tnf*, *Il1a*, and *Il1b*), and key DAM signature genes (*Apoe*, *Spp1*, *Lpl*, *Ccl3*, and *Cst7*) (Fig. 3G). Notably, CCL2/CCR2 interaction mediates the recruitment of CCR2 bearing leukocytes in the brain in several neuroinflammatory diseases (Mahad et al., 2006). Given this context, aged *Gba*^loxp/loxp^ Cx3cr1^Cre/+^ mice microglia displaying higher expression of Ccl2 and other inflammatory chemokines likely mediates infiltration of peripheral CCR2^+^ MFs and leukocytes contributing to neuroinflammation seen in our late onset neurodegeneration model, akin to late onset neuroinflammation seen in some GD patients with aging (Belarbi et al., 2020). Interestingly, expression of *Mtx1* and *Thbs3* genes that are assigned to chromosome 1q21 contiguous with *Gba*, was enriched in cluster 5, and were enriched in aged *Gba*^loxp/loxp^ Cx3cr1 microglia (we have termed these *Gba*-associated genes). The expression of IFN response genes in microglia cluster 7 remained unchanged between young and aged mice (Fig. 3E). Taken together, molecular elucidation of aged *Gba-deficient* microglia shows a loss of homeostatic signature with pronounced expression of inflammatory signaling molecules. Thus, our findings highlight the important role of *Gba* in long-term maintenance of microglial homeostasis and implicates *GBA* deficient microglia in development of age-related neurodegenerative disease like PD that occurs with high risk in adults with GD1.

**Fig. 3.**
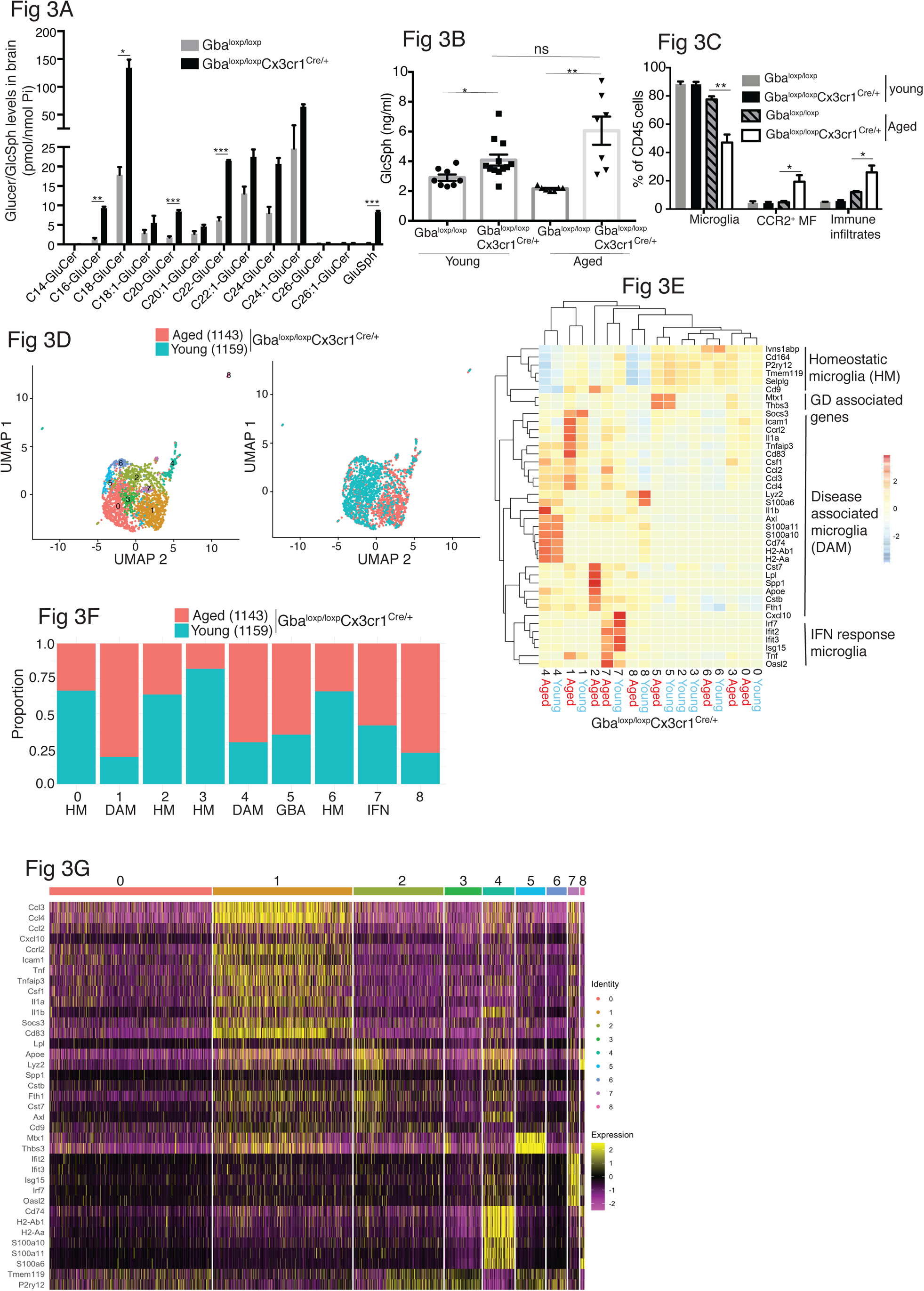
Aged Gba^loxp/loxp^ Cx3cr1^Cre/+^ mice brain show alteration of microglia subsets and neurodegeneration. **(A)** Quantitative analysis of total GluCer/GlcSph levels in Gba^loxp/loxp^ Cx3cr1^Cre/wt^ mice and control mice (n=3 mice/group). Statistical significance was determined using t test with using Bonferroni-Dunn correction for multiple comparisons (*p<0.05, **p<0.001 and ***p<0.0001) (**B)** Quantitative analysis of serum GlcSph levels in young and aged Gba^loxp/loxp^ Cx3cr1^Cre/wt^ mice and control mice (n=8-11 mice/group; repeated at least 3 times). (**C)** Bar graph shows comparison of percentage of CCR2+ MFs, microglia and immune infiltrates between young and aged Gba^loxp/loxp^ Cx3cr1^Cre/wt^ mice and control mice (n=3 mice/group; repeated at least 3 times). (**D)** UMAP plots show clustering of microglia from young and aged Gba^loxp/loxp^ Cx3cr1^Cre/wt^ mice. Cells are colored by cluster (Left) and by age (Right). (**E)** Hierarchical heat map depicting differential expression of genes taken from Wang et al and compared between microglia cluster from young and aged Gba^loxp/loxp^ Cx3cr1^Cre/wt^ mice. (**F)** Fraction of cells for each cluster present in young and aged Gba^loxp/loxp^ Cx3cr1^Cre/wt^ mice respectively. (**G)** Gene expression heat map for clusters defined as microglia. (**A-B)** Data represents 3 biological replicates. Means ± SEM are shown. Unpaired t-test, two tailed was used to test significance *p<0.05, **p<0.001 and ***p<0.0001.

### NK cells together with microglial activation drives neuropathology in nGD brains

In the characterization of immune cell infiltrates by scRNA seq in the brains of nGD and nGD Cx3cr1^Cre/+^ mice, we found evidence of infiltration of NK cells (fig. S1B and E). We confirmed striking NK infiltration in nGD brains by flow cytometry (Fig. 4 A, B, and C). Brain infiltrating NK cells expressed granzyme A (GzmA) (Fig 4 D, E, and F). NK cell frequency or GzmA expression was unaltered in the spleens of both nGD and nGD Cx3cr1^Cre/+^ mice (fig. S6A, Fig. 4G and H), implying that the primary cues responsible for activation of NK cells were emanating from within the nGD brains. Sphingosine 1-phosphate (Sph-1P) plays a key role in NK cell trafficking via its receptor S1P5 (Walzer et al., 2007). To address whether Sph-1P is involved in NK cell infiltration in nGD brain, we performed HPLC-MS/MS analysis of brain tissue for sphingosine species. We found no specific buildup of sphingosine lipid species including Sph-1P in nGD and nGD Cx3cr1^Cre/+^ mice brains compared to the control mice (fig. S6B), suggesting that other mechanism(s) may underlie NK cell infiltration or that there is local generation of S1P that is beyond the resolution of HPLC-MS/MS.

**Fig. 4.**
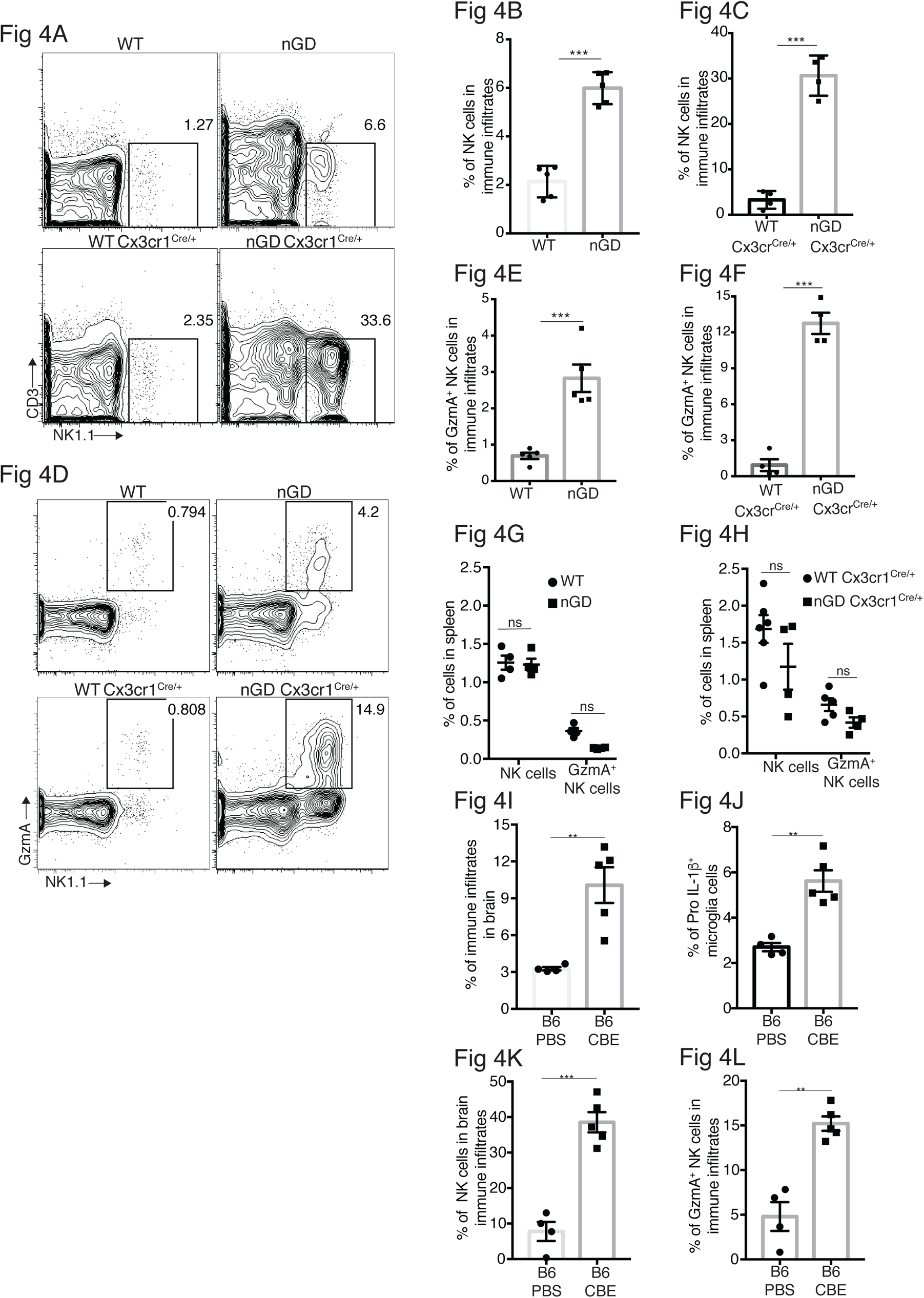
NK cell infiltration into the brain of nGD, nGD Cx3cr1^Cre/+^ mice. (**A)** CD45^+^ cells form the whole brain of nGD, nGD Cx3cr1^Cre/+^ and control mice were gated to analyze CD3^−^NK1.1^+^ NK cells. Bar graphs represent the percentage of NK cells in the immune infiltrates of (**B)** nGD mice brain (n=5 mice/group) **and (C)** nGD Cx3cr1^Cre/+^ mice brain respectively (n=4 mice/group). (**D)** Expression of granzyme A (GzmA) in the brain infiltrating NK cells of nGD and nGD Cx3cr1^Cre/+^ mice. Bar graphs compares the percentage of GzmA^+^ NK cells in in the immune infiltrates of (**E)** nGD vs control mice brain (n=5 mice/group) and **(F)** nGD Cx3cr1^Cre/+^ vs control mice brain respectively (n= 4 mice/group). Percentage of NK1.1^+^ NK cells and GzmA^+^ NK cells in the spleen of (**G)** nGD mice and (**H)** nGD Cx3cr1^Cre/+^ mice (n=4-5 mice/group). Bar graph showing percentage of (**I)** immune infiltrates (**J)** Pro-IL-1ß^+^ microglia cells (**K)** percentage of NK cells and **L.** GzmA^+^ NK cells in the whole brain of Gba^wt/wt^ treated with vehicle or CBE. Experiments were repeated thrice. (**A)** and **(D)** were representative from one of the experiments, (**B), (C), (E-L)** data are combined from 2 such experiments. Data are shown as means ± SEM. Unpaired t-test, two tailed was used to test significance. *p<0.05, **p<0.001 and ***p<0.0001.

To assess whether the pattern of neuroinflammation observed in genetic models of nGD could be replicated in another system, we used a chemically induced model of nGD using an inhibitor of acid β-glucosidase (conduritol β-epoxide, CBE). Administration of CBE in WT mice resulted in the nGD phenotype, as described previously (Farfel-Becker et al., 2011; Kanfer et al., 1982). FACS analysis of immune cells in the brains of CBE-treated mice revealed major infiltration of immune cells (fig. S6C and Fig. 4I) as well as Pro-IL-1ß induction in the microglia (Fig. 4J). Like genetic models, immune cell phenotyping in the brains of CBE-treated mice showed striking induction of GzmA^+^ NK cells (fig. S6C and Fig. 4K and L). Therefore, using genetic and chemically induced nGD mouse models, we confirmed a conserved immune landscape of nGD brains comprising prominent NK cell infiltration and microglial activation.

### Single cell resolution revealed role of microglial *Gba* in delaying neuroinflammation

To gain a comprehensive and dynamic overview of the underlying mechanisms involved in the longitudinal course of GD associated neurodegeneration, we performed brain single-nucleus RNA-seq to elucidate cell types/pathways involved in (1) the acute neuropathic model nGD mice at 2wks, (2) after restoration of *Gba* in microglia (nGD Cx3cr1^Cre/+^) both at early (2 wks) and late stage (6 wks) when it displays overt neurodegenerative phenotype, and (3) late onset model, nGD Nes^Cre/+^ brains with rescue of *Gba* in neuronal compartment (∼7mo). Integrated hierarchical analysis with quality controls (fig. S7A), revealed 61 distinct cell clusters that were annotated for major cell types based on the differential expression of canonical marker genes (Fig. 5A and fig. S7B). We focused on four distinct gene sets: lysosomal biology, ISGs, chemokines, and ApoE. Lipid metabolism genes are known to be upregulated in DAMs. We focused on ApoE because of its established roles in both lipid transport and neurodegeneration highlighted by GD (Serrano-Pozo et al., 2021). AUCell analysis was applied to interrogate these gene set enrichments and stratify different cell types: Lysosomal biology (*Abca2*, *Ap1b1*, *Ap3s1*, *Ap4s1*, *Atp6v0a1*, *Atp6v1h*, *Cd63*, *Cd68*, *Cln3*, *Ctsb*, *Ctsd*, *Ctsl*, *Ctss*, *Galc*, *Galns*, *Hexa*, *Hexb*, *Lamp*, *Psap*, *Slc17a5*, *Sumf1*, *Gpr37*, *Gpr37l1*, *Mtx1 and Thbs3*), Interferon Signaling Genes (*Psmb10*, *Psmb9*, *Psmb8*, *Psma4*, *Psme2*, *Oas1a*, *Oasl2*, *Oasl1*, *Isg15*, *Ifi207*, *Ifi47*, *Ifit1*, *Ifi213, Ifitm3*, *Ifit3*, *Ifi35*, *Epsti1*, *Parp14*, *Bst2*, *Irf7*, *Irf9*, *Stat1*, *Stat2*, *Usp18*, *Ifit2*, *Slfn5* and *Ifih1*), Chemokine genes (*Cxcl10*, *Cxcl12*, *Ccl5*, *Cxcl5*, *Ccl2*, *Ccl3* and *Ccl7*) and ApoE. In the brains of nGD and 6-week-old nGD Cx3cr1^Cre/+^ mice both with overt neurodegenerative phenotypes, lysosomal biology, ISG, and ApoE genes were highly upregulated in microglia, Purkinje, oligodendrocytes (clusters 2, 34, and 36), astrocytes (clusters 22, 23, 4, 48, 52, and 9), and neurons (0, 3, 5, and 6) (Fig. 5B). In contrast, microglia of 2-week-old nGD Cx3cr1^Cre/+^ mice were similar to controls. In line with these observations, DAM markers were significantly enriched in microglia from both nGD and 6-week-old nGD Cx3cr1^Cre/+^ compared to 2-week-old nGD Cx3cr1^Cre/+^ mice (fig. S7C), consolidating the importance of microglial *Gba* in ameliorating neurodegeneration seen in florid acute neuroinflammation. Importantly, the results suggest that even in setting of normal microglia *Gba* in nGD Cx3cr1^Cre/+^ mice, increased flux of glucosylceramide lipids arising from *Gba* deficient neuronal cells lead to DAM, over time as seen at 6 weeks but not at 2 weeks in these mice. *Gba* rescue in the neuronal compartment resulted in the normalization of pathways associated with lysosomal biology genes, ISG genes, chemokine genes, and ApoE in microglia, Purkinje, oligodendrocytes, astrocytes, and neuron cell clusters, and in other brain cell clusters (Fig. 5B). Nevertheless, lower expression of homeostatic microglia markers was observed in nGD Nes ^Cre/+^ microglia as compared to control mice (fig. S7D).

**Fig. 5.**
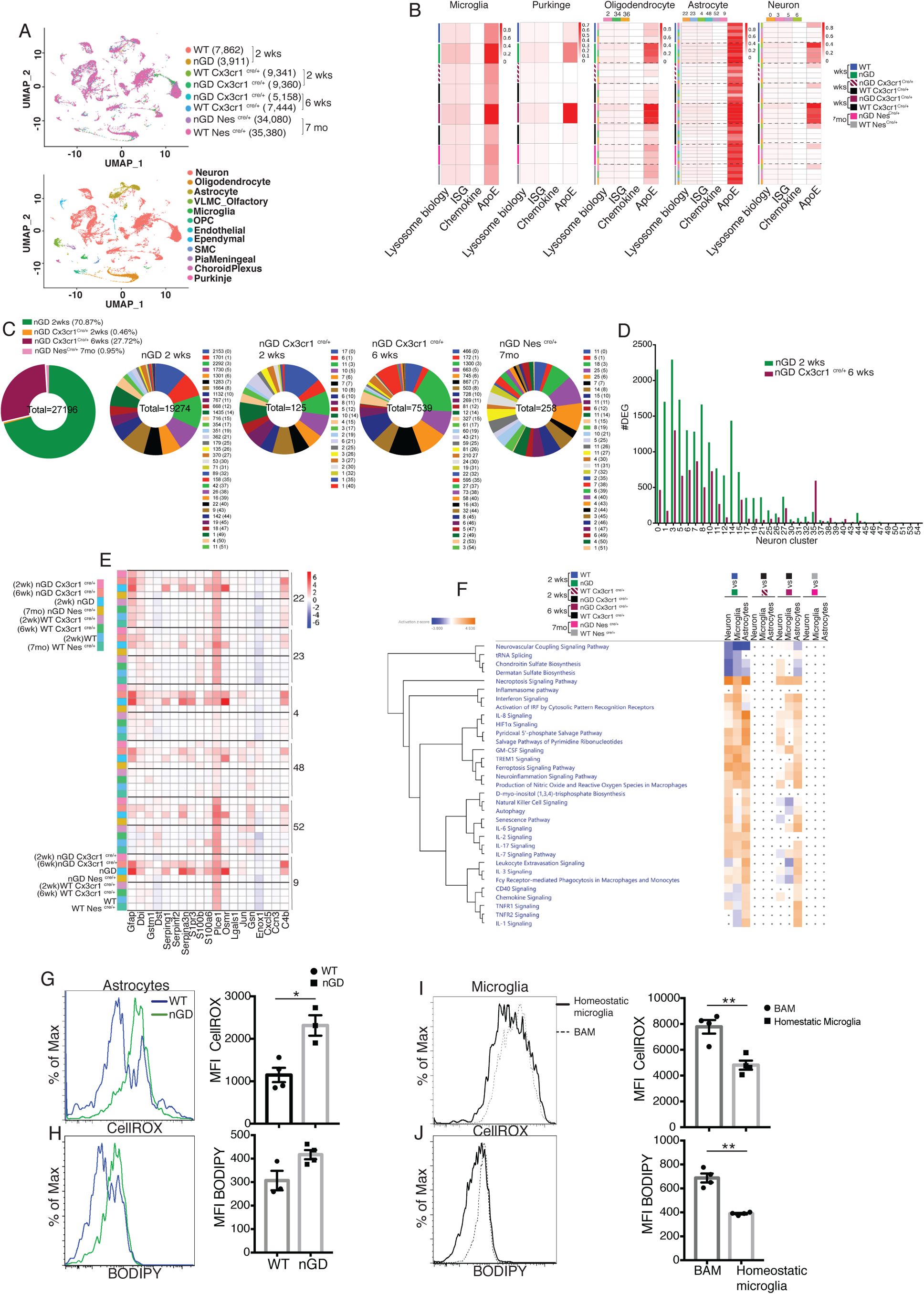
snRNAseq reveals key role of microglia astrocytes and neurons in GD associated neuroinflammation. **(A)** UMAP data from nGD (2 wk. old), nGD Cx3cr1^Cre/+^ (2 and 6 wk. old respectively), nGD Nes^Cre/+^ mice (n=3) and corresponding control mice, colored by genotype (top) and cell clusters (bottom). **(B)** AUC analysis for select lysosome, Interferon signature genes (ISG), chemokine and Apoe gene sets significantly enriched (FDR< 0.05) in microglia, Purkinje, Oligodendrocyte, Astrocyte and neuro clusters of nGD (2 wks.) vs control mice; nGD Cx3cr1^Cre/+^ (2 and 6 wks. respectively) vs control mice; nGD Nes^Cre/+^ mice vs control mice. Row side bar colors indicate mice genotype, age and cell clusters. (**C)** Pie chart displays the number of differentially expressed genes (DEGs) in neuronal clusters of nGD (2 wks.) vs control mice (green); nGD Cx3cr1^Cre/+^ (2 wks. (orange) and 6 wks. (maroon) respectively); nGD Nes^Cre/+^ mice (pink) vs control mice with log2(Fold Change) and adjusted p-value < 0.05. Total DEGs from each set of mice are stated in the middle of pie chart with number of DEGs and the neuronal cluster in brackets are shown to the right of each pie chart. (**D)** Bar graph represents the number of DEGs in neuronal clusters of nGD 2 wks. (green) vs nGD Cx3cr1^Cre/+^ 6 wks. (maroon) with log2(Fold Change) and adjusted p-value < 0.05. (**E)** Heat map of differentially expressed genes associated with Disease Associated Astrocytes (DAA) from nGD; nGD Cx3cr1^Cre/+^ (2 wks. and 6 wks. respectively); nGD Nes^Cre/+^ mice vs control mice. p<0.05 was considered significant (2-sided t tests). All individual DAA genes with significant differential expression are listed on bottom and the astrocyte clusters are shown in right. Red, positive z-score; white, zero z-score; blue, negative z-score. (**F)** Ingenuity Pathway Analysis (IPA) from nGD vs WT, nGD Cx3cr1^Cre/+^ vs WT Cx3cr1^Cre/+^ at 2 and 6 wks. old mice; nGD Nes^Cre/+^ mice vs WT Nes^Cre/+^ in neuron cluster, in microglia and Astrocytes. Orange, positive z-score; white, zero z-score; blue, negative z-score; gray dots are statistically insignificant. (**G)** Representative flow cytometry histogram (left) and quantification of CellROX fluorescence in astrocytes. (**H)** Histogram (left) and quantification (right) of BODIPY fluorescence in astrocytes of nGD mice. (**I)** Flow cytometry histogram (left) and quantification (right) of CellROX fluorescence in activated and homeostatic microglia from nGD mice. (**J)** Histogram (left) and quantification (right) of BODIPY fluorescence in activated and homeostatic microglia from nGD mice n=3-4 mice per group. Data were replicated in at least two independent experiments. Unpaired t-test, two tailed was used to test significance. *p<0.05, **p<0.001 and ***p<0.0001.

Gene expression analysis revealed that nGD mice exhibited the largest number of differentially expressed genes (DEGs) (n = 19,274), which were strikingly reduced in 2-week-old nGD Cx3cr1^Cre/+^ (n=125) and also in nGD Nes^Cre/+^ brains, n=256 (Fig. 5C). Consistent with the neuroinflammation observed in 6-week-old nGD Cx3cr1^Cre/+^ mice, elevated DEGs in neuronal clusters were seen (n=7539) compared to 2-week-old nGD Cx3cr1^Cre/+^ mice (n=125) (Fig. 5C). However, 6-week-old nGD Cx3cr1^Cre/+^ mice show a reduction of DEGs in different neuronal clusters as compared to nGD mice (Fig. 5D). Collectively, these results indicate the underlying role of microglial *Gba* in safeguarding and delaying neuroinflammatory gene networks in nGD brains.

We found evidence of significant activation of severe disease pathways in astrocytes (Fig. 5B). Astrocytosis has been described in immunohistochemistry of mouse and human neuronopathic brains (Wong et al., 2004). We analyzed the astrocyte compartment for the disease-associated astrocyte (DAA) gene signature (Batiuk et al., 2020; Habib et al., 2020). DAA genes were highly induced in astrocyte clusters from nGD mice and in 6-week-old nGD Cx3cr1^Cre/+^ mice but not in 2-week-old nGD Cx3cr1^Cre/+^ mice (Fig. 5E). IPA analysis was performed on neurons, astrocytes, and microglia. IPA revealed enrichment of necroptosis, interferon, IL8, TREM1, and neuroinflammation signaling pathways in 2-week-old nGD mice compared to the controls. *Gba* rescue in microglia of 2-week-old nGD Cx3cr1^Cre/+^ mice showed abrogation of neuroinflammatory pathways seen in nGD brains (Fig. 5F). Between 2 and 6 weeks, as neurodegeneration advanced in nGD Cx3cr1^Cre/+^ mice, the enrichment of inflammatory pathways was again evident (Fig. 5F). Notably, both interferon and NK cell signaling pathways were upregulated in 2-week-old nGD and in 6-week-old nGD Cx3cr1^Cre/+^ brains (Fig. 5F). Consistent with the lack of microglial activation and NK infiltration in nGD Nes^Cre/+^ mice, IPA pathway analysis revealed no changes in neurons, astrocytes, and microglia in nGD Nes^Cre/+^ mice (Fig. 5F). Pathway analysis of differentially expressed genes revealed that the ‘production of nitric oxide and ROS’ was elevated in nGD and 6-week-old nGD Cx3cr1^Cre/+^ microglia and astrocytes along with neuroinflammation pathways. Elevated and dysregulated reactive oxygen species (ROS) production from DAM contributes to oxidative stress and has been shown to be intricately linked with neurodegeneration (Mendiola et al., 2020; Simpson and Oliver, 2020). Consistent with the transcriptional changes observed in nGD mice, astrocytes from nGD mice showed higher fluorescence after treatment with CellROX, indicating a higher level of ROS, as compared to astrocytes from control mice (Fig. 5G). nGD astrocytes with high ROS level also showed elevated accumulation of neutral lipids, as seen by BODIPY staining (Fig. 5H). Moreover, activated microglia in the nGD brain showed enhanced ROS level along with an increased accumulation of lipids compared to homeostatic microglia (Fig. 5I and J). Overall, these observations suggest that *Gba-deficient* neurons and microglia play key roles in orchestrating neuroinflammation involving astrocytes and NK cells that underlie neurodegeneration in neuronopathic GD.

### GCS inhibitor reduces GluCer/GlcSph and reverses microglia and NK cell activation

Collectively our findings suggest that restoring lipid dyshomeostasis and neurodegeneration caused by *Gba* deficiency in microglia and neurons can be prevented by restoration of Gba. Previous studies have shown that treatment with GCS inhibitor GZ-161 reduced GluCer and GlcSph in the brains of nGD mice and prolonged survival by a few days (Cabrera-Salazar et al., 2012) (fig. S8A). To determine whether reduction of glucosylceramide lipids by GZ-161 alleviates neuroinflammatory landscape and neurodegeneration, we treated our mouse models with this brain permeant GCS inhibitor. Consistent with previous studies, GZ-161 treatment significantly prolonged the survival of nGD mice (Fig. 6A). We investigated whether GZ-161 treatment could further extend the survival of nGD Cx3cr1^Cre/+^ mice by more effective reduction of brain GluCer/GlcSph via dual effect of decreased synthesis of GluCer and increased lysosomal degradation of GluCer in microglia. Indeed, GZ-161 treatment of nGD Cx3cr1^Cre/+^ mice resulted in considerable extension of survival compared to untreated mice, with analogous normalization of GlcSph levels in the sera of treated nGD Cx3cr1^Cre/+^ mice (Fig. 6A and B).

**Fig. 6.**
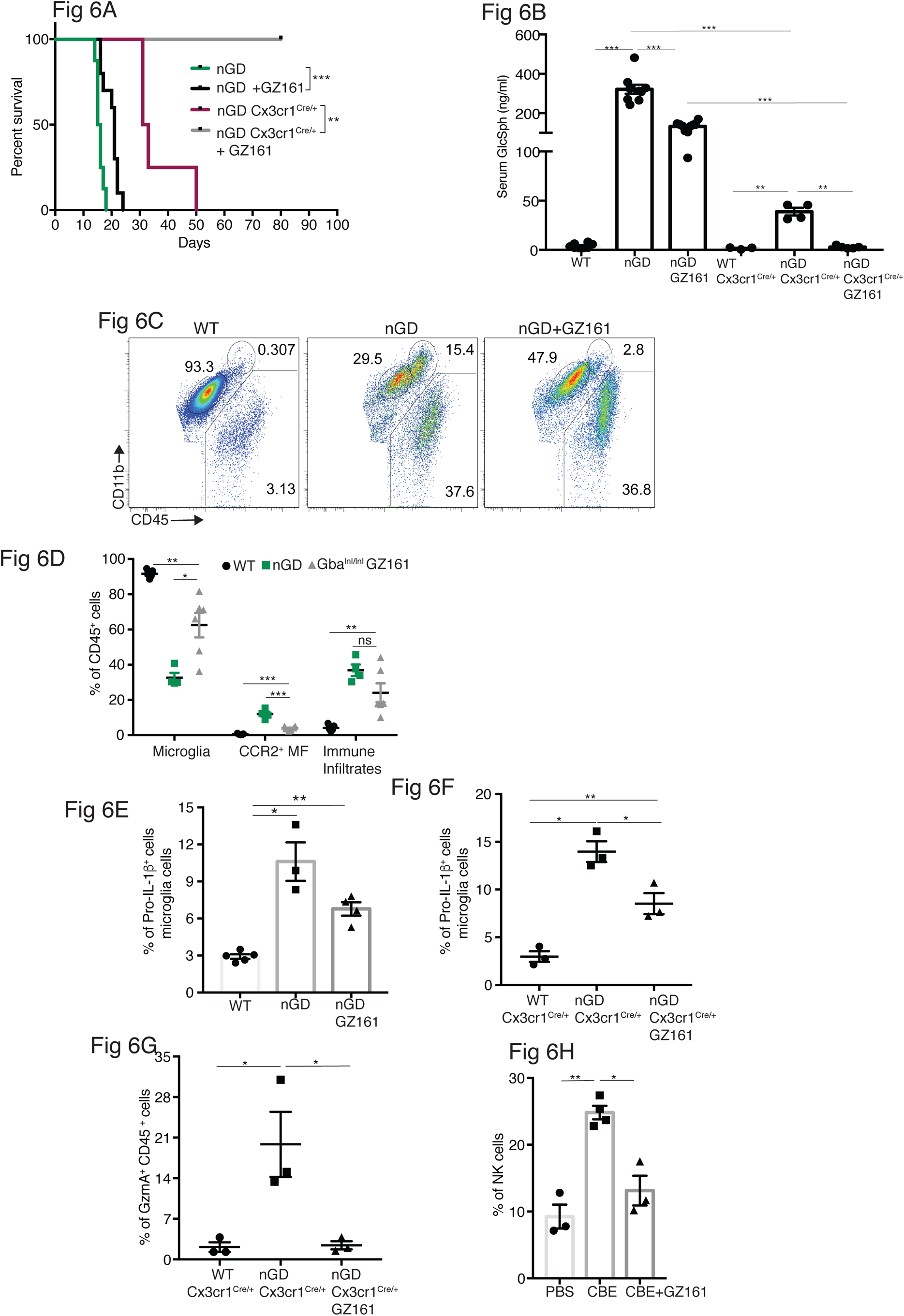
Effects of GCS inhibitor (GZ-161) on microglia and GzmA^+^ cells. **(A)** Kaplan-Meier Survival analysis of nGD and nGD Cx3cr1^Cre/+^ mice after GZ-161 treatment compared to vehicle treated controls (n=4-10 mice/group). (**B)** Quantitative analysis of serum GlcSph levels by LC-ESI-MS/ MS in nGD, nGD Cx3cr1^Cre/+^ mice with and without GZ-161 treatment compared with the vehicle treated controls (n=4-10 mice/group). (**C)** Representative FACS staining of microglia, activated microglia and immune infiltrates in wild type and nGD mice with and without treatment with GZ-161. (**D)** Graph represents percentages of microglia, CCR2+ MFs and immune infiltrates in wild type and nGD mice with and without treatment with GZ-161 (n=4-6 mice/group). (**E)** Bar graph showing percentage of Pro-IL-1ß^+^ microglia cells in wild type and nGD mice with and without treatment with GZ-161 (n=3-5 mice/group; repeated at least 3 times). **(F)** Bar graph showing percentage of Pro-IL-1ß^+^ microglia cells in wild type and nGD Cx3cr1^Cre/+^mice with and without treatment with GZ-161(n=3-5 mice/group). (**G).** Percentage of GzmA^+^ CD45^+^ cells in wild type and nGD Cx3cr1^Cre/+^mice with and without treatment with GZ-161 (n=3-4 mice/group (n=3-5 mice/group). (**H)** Bar graphs compares the percentage of NK cells in Gba^wt/wt^ and mice treated with CBE and CBE+ GZ-161 respectively (n=3-4 mice/group (n=3-5 mice/group; repeated at least 3 times). Data represents 3 biological replicates. Means ± SEM are shown. Unpaired t-test, two tailed was used to test significance. *p<0.05, **p<0.001 and ***p<0.0001.

Next, we assessed the impact of pharmacologic reduction of GluCer/GlcSph on disease-specific, microglial and macrophage phenotypes associated with neuroinflammation in GD mice models. FACS analysis of GZ-161-treated nGD mice showed that the GCS inhibitor significantly reduced the proportion of CCR2^+^ MFs, concomitantly restoring homeostatic microglia compartment (Fig. 6C and D). However, GZ-161 treatment had no effect on immune cell infiltrates in the brains of nGD mice (Fig. 6C and D). GZ-161treatment showed a non-significant downward trend in microglial Pro-IL-1ß induction in nGD mice (Fig. 6E). GZ-161 treatment in longer-lived nGD Cx3cr1^Cre/+^ mice wherein drug administration was more reliable resulted in significantly increased survival (Fig. 6A) accompanied by a reduction in Pro-IL-1ß^+^ microglia as well as a reduction in GzmA^+^ immune cells in the brain (fig. S8B, Fig. 6F and G). We further evaluated the ability of GZ-161 to counteract NK cell induction seen in the CBE model. Administration of GZ-161 in CBE-treated mice reduced infiltration of NK cells in the brain (Fig. 6H). Together, these results show that GZ-161 can mitigate the metabolic defect caused by *Gba* deficiency by reducing the toxic accumulation of GluCer and GlcSph, concurrent with alleviating CNS immune inflammation involving microglia and NK cell activation. Moreover, when microglia are predominantly of homeostatic phenotype through complementation of *Gba* function in nGD mice, a two-pronged approach of restoring *Gba* in microglia and GCS inhibitor, the influx of NK cells into the brain and subsequent neurodegeneration is significantly constrained.

### GCS inhibition counteracts age-related microglial dysfunction and NK cell activation

We investigated whether buildup of GlcCer and GlcSph is proximate cause of neuroinflammation and neurodegeneration in *Gba* deficiency using the brain permeant GCS inhibitor in late onset neurodegeneration *Gba*^loxp/loxp^ Cx3cr1^Cre/+^ model. The effect of reversing the toxic accumulation of GluCer/GlcSph on *in vivo* microglial homeostasis and NK cell activation was assessed by feeding *Gba*^loxp/loxp^ control mice and *Gba*^loxp/loxp^ Cx3cr1^Cre/+^ mice with either control or GZ-161-formulated diet, starting at age 3-months, for 7 months. The effect of GZ-161 on GSL accumulation in brain regions was evaluated using MALDI imaging. Gba^loxp/loxp^ Cx3cr1^Cre/+^ mice accumulated HexCer (d18:1/18:0) and HexCer (d18:1/20:0) in the corpus callosum (Fig. 7A), which was normalized upon GZ-161 diet. Concomitantly, there was a reduction in HexCer (18:1/22:0) accumulation in the cerebral cortex (Fig. 7A). Interestingly, as previously stated, LysoPC was present with an exceptionally strong signal in the microglia in the cerebral cortex and midbrain regions of untreated *Gba*^loxp/loxp^ Cx3cr1 ^Cre/+^ mice. Intensity of LysoPC accumulation in brain microglia was significantly reduced by GZ-161 treatment.

**Fig. 7.**
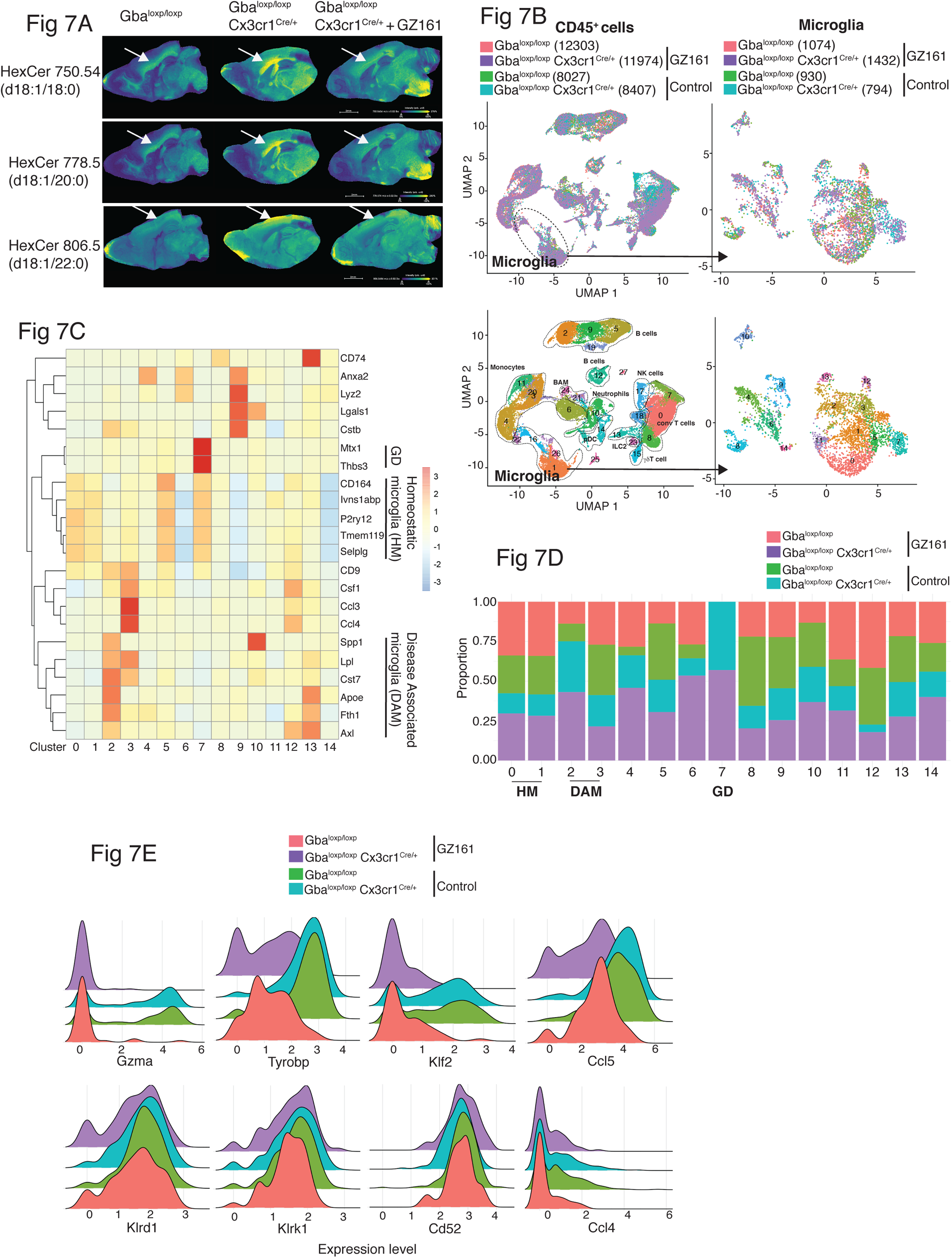
Long term treatment with GCS inhibitor, GZ-161, counteracts age-related microglial dysfunction and NK cell activation. **(A)** Signal intensities of HexCer species and LysoPC identified by MALDI across Gba^loxp/loxp^ Cx3cr1^Cre/wt^ mice treated with either vehicle or GZ161 and control mice brain. The color bars in MALDi images show signal intensity: blue to yellow indicates low to high levels. (**B)** UMAP plots show clusters of CD45^+^ cells from brain of control and Gba^loxp/loxp^ Cx3cr1^Cre/wt^ mice treated with either GZ161 or vehicle. Cells are colored by mice genotype (Left top) and by cluster (Left bottom). Microglia sub cluster of CD45^+^ cells, colored by mice (Right top) and by clusters (Right bottom). (**D)** Hierarchical heat map depicting differential expression of genes associated with homeostatic and disease associated microglia in the different microglia clusters. (**D)** Fraction of cells present in each microglial cluster from control and Gba^loxp/loxp^ Cx3cr1^Cre/wt^ mice treated with either GZ161 or vehicle. (**E)** Histogram showing differential expression of selected genes in cluster 17 from control and Gba^loxp/loxp^ Cx3cr1^Cre/wt^ mice treated with either GZ161 or vehicle.

We performed scRNA-seq of CD45^+^ cells from brains of *Gba*^loxp/loxp^ Cx3cr1^Cre/+^ mice and control mice on control or GZ-161-formulated diet. A total of 40,763 CD45^+^ single-cell transcriptomes were subjected to unsupervised Louvain clustering resulting in a total of 28 transcriptionally distinct populations comprising 10 broad cell types (Fig. 7B). Clusters 1, 26, and 16 represented microglia (4,230 cells) and were visualized on UMAP, revealing 15 unique clusters (Fig. 7B, lower panel). We identified clusters 0, 1, and 5 as homeostatic microglia, cluster 2 as DAM, and cluster 7 as *Gba*-associated microglia (Fig. 7C). DAM cluster 2, which was significantly enriched in *Gba*^loxp/loxp^ Cx3cr1^Cre/+^ brains was markedly reduced in *Gba*^loxp/loxp^ Cx3cr1^Cre/+^ mice fed with GZ-161 diet with concurrent enhancement of homeostatic microglia clusters 0 and 1 (Fig. 7D). We found that *Gba-associated* gene cluster 7 was present exclusively in *Gba*^loxp/loxp^ Cx3cr1^Cre/+^ mice both on GZ-161 and on the control diet groups. To assess the effect of GZ-161 on NK cells within the brain environment, we further analyzed cluster 17, which represented NK cells (fig. S9A). GZ-161 treatment resulted in a reduction of *Gzma*, *Ccl5*, *Tyrobp*, and *Klf2*, with no change in *Klrd1, Klrk1, CD52*, and *Ccl4* (Fig. 7E). Collectively, these results demonstrate that GCS inhibitor-mediated reduction of GluCer and GlcSph counteracts age-related microglial dysfunction and NK cell activation in several brain regions in *Gba*^loxp/loxp^ Cx3cr1 ^Cre/+^ mice. Together, these findings not only validate GlcCer/GlcSph as the proximate cause of neuroinflammation involving microglia and NK cells but also promising utility of brain permeant small molecule inhibitor as a therapeutic approach for neuronopathic GD.

### Generating novel candidate biomarkers of neurodegeneration in *Gba*-deficiency

Given the striking relationship between serum Nf-L, level of pathogenic lipid, GlcSph, and neurodegenerative phenotype, we sought to validate these biomarkers in translational studies in GD patients and explore other potential biomarkers suggested by our results. Lack of circulating biomarkers for pre-symptomatic *Gba* deficiency associated neurodegenerative diseases is a major impediment to optimal management of patients and conduct of clinical trials. Therefore, we investigated if serum levels of neurofilament light (Nf-L), an accepted biomarker of neuroaxonal injury in several neurodegenerative and neuroinflammatory diseases (Gaetani et al., 2019; Khalil et al., 2020; Loeffler et al., 2020; Weinhofer et al., 2021), could serve as a biomarker of neurodegeneration in nGD models. Using ultra-sensitive Quanterix Simoa™, we found a massive 2,000-fold elevation of serum Nf-L in nGD mice, ∼100-fold elevation in nGD Cx3cr1^Cre/+^, and ∼20-fold elevation in nGD Nes^Cre/+^ mice (Fig. 8A). Remarkably, GZ-161 treatment which led to reductions in GlcSph levels also led to significant decline in serum Nf-L levels in nGD and nGD Cx3cr1^Cre/+^ mice (Fig. 8A). Serum Nf-L level in nGD mice (18,100± 4809 pg/ml) was reduced by GZ-161 treatment to 6,876±1080 pg/ml. Conversely, GZ-161 treatment of nGD Cx3cr1^Cre/+^ mice dramatically reduced Nf-L level to 1297±534.1 pg/ml compared to untreated mice (5466±557.2 pg/ml). The residual Nf-L level after GZ-161 treatment in nGD Cx3cr1^Cre/+^ mice was significantly lower than that in nGD mice treated with GZ-161, consistent with the synergistic effect of GCS inhibition and restoration of *Gba* function in microglia (Fig. 8A). Increased survival of nGD and nGD Cx3cr1^Cre/+^ mice after GZ-161 treatment correlated with amelioration of neurodegeneration, as indicated by serum Nf-L levels. Notably, consistent with the age-related progression of neuroaxonal damage, there was a clear age-related increase in serum Nf-L levels with the most significant increase in aged *Gba*^loxp/loxp^ Cx3cr1^Cre/+^ mice (Fig. 8B). Serum levels of Nf-L and GlcSph were strongly correlated (r=0.8344, p<0.0001), consistent with a direct link between lysosomal generation of toxic GlcSph and neuroaxonal injury (Fig. 8C). Together, these data support the notion that GlcSph (and GluCer) reduction therapy with the brain-penetrant GCS inhibitor GZ-161 ameliorates neurodegeneration in nGD mice. Moreover, the results suggest that serum Nf-L and GlcSph are promising biomarkers of neurodegeneration in *Gba* deficiency.

**Fig. 8.**
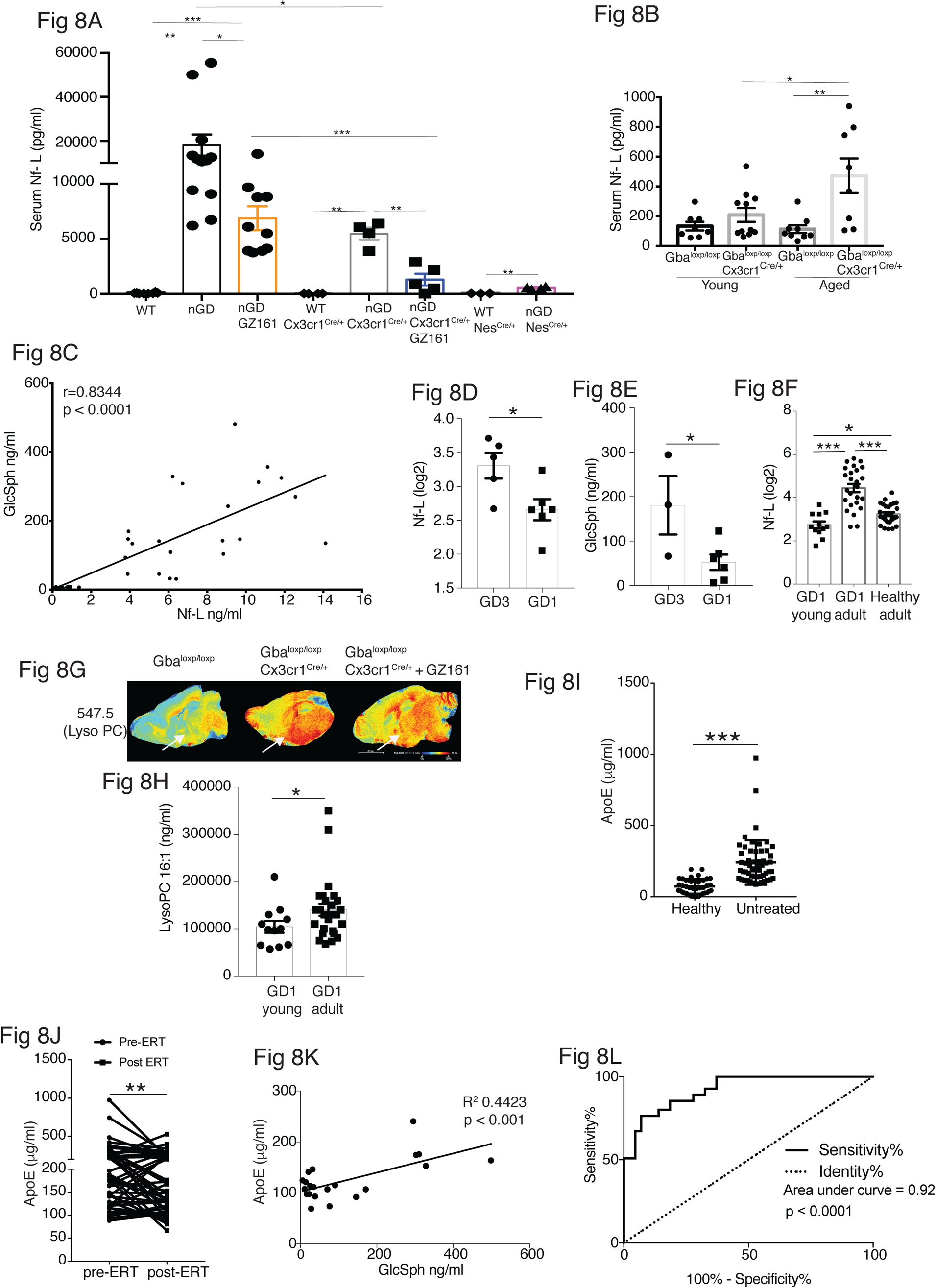
Clinical evaluation of ApoE and Nf-L as biomarkers of GD associated neurodegeneration. **(A)** Quantitative analysis of serum Nf-L levels in nGD, nGD Cx3cr1^Cre/+^ and nGD Nes^Cre/+^ mice with and without GZ-161 treatment compared with the vehicle treated controls (n=4-10 mice/group; two independent experiments). (**B)** Comparison of serum Nf-L levels between young and aged Gba^loxp/loxp^ Cx3cr1^Cre/wt^ mice along with control mice (n=8-11 mice/group). (**C)** Correlation between serum GlcSph and Nf-L levels in nGD, nGD Cx3cr1^Cre/+^ mice, nGD Nes^Cre/+^ and control mice. The p value obtained from Spearman’s rank correlation coefficient test was < 0.0001 (n=69 mice). (**D)** Quantitative analysis of serum Nf-L levels (log2 scale) in GD 3 patients (n=5) compared with age matched GD1 patient (n= 6). (**E)** Quantitative analysis of serum levels GlcSph in GD 3 patients (n=3) compared with age matched GD1 patient (n= 6). (**F)** Quantitative analysis of serum Nf-L levels (log2 scale) in young GD1 patients (n=12) compared with adult GD1 patient (n= 25) and adult healthy controls (n= 28). (**G)** Signal intensities of LysoPC identified by MALDI across Gba^loxp/loxp^ Cx3cr1^Cre/wt^ mice treated with either vehicle or GZ161 and control mice brain. The color bars in MALDi images show signal intensity: blue to red indicates low to high levels. **(H)** Quantitative analysis of serum levels of LysoPC 16:1 species in young GD 1 patients (n=12) compared with adult GD1 patient (n= 26). (**I)** Graph represents Apoe levels in the sera of untreated GD1 patients (n=55) and healthy controls (n=43). (**J)** Graph represents Apoe levels in the sera of untreated GD1 patients (n=55) and after Enzyme Replacement Therapy (ERT) (n=55). (**K)** Correlation between serum GlcSph and ApoE levels in sera of GD1 patients. The p value obtained from Pearson’s correlation test was < 0.001 (n=21 patients). (**L)** ROC curves for serum ApoE expression in GD1 patients and Area Under the Curve (AUC). Means ± SEM are shown. Differences between groups were analyzed using unpaired t-test A-B Mann-Whitey test (C,E,H and I), two tailed. *p<0.05, **p<0.001 and ***p<0.0001.

Our findings of serum Nf-L correlating with severity of neurodegeneration as well as serum levels of GlcSph in various permutations of nGD mice model, for the first time reveal that such circulating biomarkers may exist not only for tracking the severity of neurodegeneration but also for monitoring the response to therapy. Therefore, we conducted an analysis of several candidate biomarkers of neurodegeneration in patients with GD. We compared serum Nf-L levels in GD3 patients with early mild neurodegenerative symptoms with age-matched GD1 patients who did not develop early onset neurodegeneration. Serum Nf-L levels were elevated in GD3 patients compared to those in GD1 (Fig. 8D). This increase in serum Nf-L levels was associated with elevated serum GlcSph levels in GD3 patients (Fig. 8E). Akin to age-related increases in serum Nf-L levels in *Gba*^loxp/loxp^ Cx3cr1^Cre/+^ mice, indicative of age-related progression of neuroaxonal damage (Fig. 8B), serum Nf-L levels were significantly elevated in adult GD1 patients as compared to children with GD1 and adult healthy controls (Fig. 8F). This is significant, as adult GD1 patients are at an increased risk of Parkinson’s disease/Lewy body dementia. The finding of striking LysoPC accumulation in *Gba*-deficient microglia (Fig. 8G) prompted us to measure LysoPC in the sera of GD1 patients, comparing young and older patients. Similar to the results with serum Nf-L, older GD1 patients exhibited increased serum levels of LysoPC 16:1 (Fig. 8H). These findings raise the possibility of broader involvement of lipid metabolism pathways beyond the primary *Gba* defect in GD and fuller understanding could lead to useful biomarkers for the clinic and role of co-regulated lipids in pathophysiology that may be targeted by treatments for GD.

Next, building upon the striking induction of ApoE expression beyond the astrocytes seen in our study, we examined ApoE as a potential biomarker for GD. In steady state, ApoE plays a critical role in lipid metabolism in the brain, with the largest contribution from astrocytes. In disease settings, the induction of ApoE is among the key signature genes of DAM. Thus, the role of ApoE in the brain is multifaceted, including lipid homeostasis, modulation of neuroinflammation, and neuronal repair (Serrano-Pozo et al., 2021). Hence, in the context of an inborn error of lipid metabolism exemplified by Gaucher disease with prominent neurodegeneration, it is highly relevant to explore ApoE expression in various brain cells in different states of nGD. We found prominent ApoE expression in astrocytes and activated microglia in nGD mice. Additionally, there was a striking induction of ApoE in multiple neuronal cell types as well as in endothelial cells (fig. S9B). Interestingly, there was concomitant induction of *ABCA1*, suggesting coupling with increased efflux of accumulating pathogenic lipids from cells in *Gba* deficient brain (fig. S9C). Therefore, we measured ApoE in the sera of 55 adults with confirmed GD1 before and after enzyme replacement therapy (ERT), which dramatically lowered GlcSph and reversed systemic clinical manifestations. Sera of untreated GD1 patients showed markedly increased levels of ApoE compared to healthy control sera (Fig. 8I). ERT resulted in a marked decrease in ApoE levels (Fig. 8J). Notably, elevated ApoE levels correlated significantly with serum GlcSph levels (Fig. 8K). The sensitivity and specificity of ApoE as a biomarker were underscored by the area under the receiver operator characteristics (ROC) of 0.92 (*p* < 0.001) (Fig. 8L), similar to that seen in other accepted biomarkers of GD, such as chitotriosidase, gpNMB and GlcSph (Murugesan et al., 2016).

## DISCUSSION

There is limited information of how *Gba* deficiency affects the cross talk between immune cells and neuronal cell types due to accumulating glucosylceramide lipids. Hence, there are no effective therapies for devastating neurodegenerative diseases associated with *Gba* mutations, and clinical trials have been severely hindered by a lack of reliable biomarkers. Previous studies in mice have suggested utility of brain permeant GCS inhibitor as treatment for neuronopathic GD but a clinical trial of Miglustat (N-butyldeoxynojirimycin), an iminosugar did not improve neurological end-points in GD3(Schiffmann et al., 2008). Subsequently, more potent GCS inhibitors based on ceramide analogs have shown more promise in genetic nGD model and in chemically induced model (Blumenreich et al., 2021; Cabrera-Salazar et al., 2012; Shayman, 2010; Wilson et al., 2020).

To delineate the impact of glycosphingolipid accumulation, identify candidate biomarkers, and assess the impact of GluCer/GlcSph lowering using a potent inhibitor of GCS, we developed an array of genetic mouse models to probe the cell-specific roles of *Gba* at single-cell resolution. Our models recapitulated both early as well as late-onset neurodegenerative GD, that provided unique insights through resolving the heterogeneity of brain macrophage populations. Our findings implicate the essential role of GlcCer-laden microglia and immune infiltrates (including CCR2^+^ MFs defined as CD11b^hi^ CD45^+^CCR2^+^ CD64^+^ TIMD4^−^ population and NK cells)and astrocytes in orchestrating neuroinflammation. Our studies identified key attributes of GD associated neuroinflammation in the form of attrition of homeostatic microglia, emergence of DAM, influx of CCR2^+^ MFs, activation of the ISG pathway and infiltration of activated NK cells. Massive cellular accumulation of GluCer/GlcSph due to *Gba* deficiency in microglia, immune infiltrates and neurons resulted in early onset of neuroinflammation which was attenuated into late-onset neurodegenerative disease by selective rescue of *Gba* in either microglia or neurons as well as by pharmacological reduction of GluCer/GlcSph in the brain using GCS inhibitor. Interestingly, inflammatory microglia also expressed several proposed biomarkers of GD, that we and others have reported such as complement pathway genes, gpNMB, cathepsin D, and cathepsin S (Afinogenova et al., 2019; Mistry et al., 2014; Murugesan et al., 2016; Pandey et al., 2017). Rescue of *Gba* in neuronal progenitors increased survival, but nevertheless, markedly shortened compared to control mice, requiring humane endpoint sacrifice due to morbid conditions and autistic behavior, reminiscent of human GD3 (Abdelwahab et al., 2017; Bilbo and Stevens, 2017; Wong et al., 2004). Late-onset neurodegenerative disease observed in some patients with GD1 (Bultron et al., 2010) appeared to be recapitulated in *Gba*^loxp/loxp^ Cx3cr1^Cre/+^ mice having slow progressive neuroinflammation with accumulation of GluCer lipids in the brain, elevated serum Nf-L and DAM gene signatures.

In our nGD mouse models with robust neuroinflammation we found prominent GzmA^+^ NK cell infiltration. Restoring *Gba* function in microglia alone showed limited effect on NK cell infiltration, however treatment with GCS inhibitor not only offset GluCer/GlcSph buildup, but also exhibited a novel immunomodulatory effect by abrogating NK cell activation. Therefore, combination therapy with GCS inhibitors and microglia-targeted therapeutics (Xu et al., 2020) merits further exploration. In general, the immune response of NK cells is fine-tuned by a balance of stimulatory and inhibitory signals based on a distinct receptor repertoire. In certain pathological states, injured neurons display activating NKG2D ligands that target them for NK cell-mediated injury(Davies et al., 2019). A potential mechanism relevant to our findings in nGD, for NK cell involvement in neurodegeneration is via HLA-1 recognition by NK cells through its inhibitory receptors Ly49 (in mice) and KIR (in humans), which mediate self/non-self-discrimination (Colonna and Samaridis, 1995; Karlhofer et al., 1992). Thus, downregulation of HLA-1 surface expression is envisioned to trigger NK cell-mediated neuronal injury due to ‘missing self’. Consistent with this notion, a recent study reported that altered cell surface GSL repertoire limited accessibility of HLA-1 by immune cells, such as CD8 T cells and NK cells, and treatment with a GCS inhibitor (N-butyl-deoxynojirimycin) fully restored accessibility to HLA-1 (Jongsma et al., 2021). Indeed, in our nGD models, a vastly more potent GCS inhibitor, GZ-161 was highly effective in reversing NK cell activation concomitant with marked reduction of brain glucosylceramide lipids. Therefore, it seems likely that in severe *Gba* deficiency, causing early onset neurodegeneration, altered cell surface GSLs in neurons impair the accessibility of HLA-1 by NK cell receptors, triggering activation and neuronal degeneration.

Our studies show remarkable efficacy of potent brain penetrant GCS inhibitor, GZ-161 in reducing GluCer and GlcSph in immune and neuronal cells of nGD mice concomitant with amelioration of neuroinflammation and neurodegeneration. A prior clinical trial of GCS inhibitor N-butyldeoxynojirimycin in GD3 showed no effect on neurological symptoms(Schiffmann et al., 2008). In such clinical trials patients have established advanced neurological disease which can hinder assessment of full therapeutic effects. Therefore, a major unmet need in *GBA1* mutation associated neurodegenerative disease is the lack of suitable biomarkers to detect neurodegeneration before onset of overt neurological symptoms, such as saccades, ataxia, or seizures. We aimed to leverage the findings from our nGD models to generate novel biomarkers. In our study, neuroaxonal injury was reflected in the elevation of serum Nf-L (Gaetani et al., 2019; Weinhofer et al., 2021). Data from our nGD models with different rates of neurodegeneration, revealed Nf-L as a strong serum biomarker of neurodegeneration that correlated with survival as well as with serum GlcSph. The candidacy of these biomarkers is especially bolstered by the finding of rising levels of serum Nf-L and GlcSph in *Gba*^loxp/loxp^ Cx3cr1^Cre/+^ mice, as the only source of these biomarkers in these mice is from within the brain. These findings raise exciting opportunities to explore serum Nf-L as a biomarker of neurodegeneration to help address some major challenges in the management of patients with GD, that is, distinguish between GD2 and early onset GD3, track subclinical neurodegeneration in the extremely variable GD3, to help individualize future therapies and to identify GD1 individuals at-risk for PD/LBD. Indeed, we found that serum Nf-L levels were higher in GD3 patients than in age-matched GD1 patients. Moreover, Nf-L levels were higher in older patients with GD1 compared to younger patients. This is significant considering that older GD1 patients are at a risk of developing PD/LBD. In this context, our findings of serum Nf-L as a strong biomarker raise the exciting prospect of detecting subclinical neurodegeneration, allowing early start of treatment to achieve better clinical outcome. There is an emerging consensus that providing treatment before the onset of overt disease manifestations offers prospects for best outcomes in inborn errors of metabolism. Studying larger cohorts of patients stratified by different types of neurodegeneration due to Gba mutations may aid further biomarker validation. The second promising biomarker emanates from striking induction of ApoE in all neuronal cell types beyond astrocytes and DAM in nGD models. Our translational result of striking elevation of ApoE levels in GD patients that correlates with GlcSph levels is compelling to explore the role of ApoE in neurodegeneration associated with Gba mutations. Indeed, a recent study showed increased prevalence of ApoE4 allele in heterozygote carriers of *GBA1* mutations who develop PD (Shiner et al., 2021).

Overall, through our systematic single cell transcriptome analysis, we identified cell populations, immunological and pathophysiological mechanisms underlying neurodegeneration associated with *GBA1* deficiency. This approach also yielded therapeutic targets and highly promising biomarkers for clinical validation to improve patient care and aid clinical trials.

### Limitations of the study

There are several limitations of our studies. In Gba^lnl/lnl^ model of fulminant neurodegeneration that phenocopies human GD2, GCS inhibitor prolonged survival significantly but the overall effect was modest. This is likely related to challenges in administering GCS inhibitor by gavage in the first days of life. While our study showed a critical role for both neuronal and microglial *Gba* in Gaucher-related neurodegeneration and involvement of NK cells, it was beyond the scope of current work to delineate a hierarchy of neuronal populations according to their vulnerability to toxic accumulation of lipids in setting of *Gba* deficiency and compensatory or pathological interactions with glial cells, infiltrating NK cells and other immune cells. With regard to application of our findings to human health, an earlier randomized trial of GCS inhibitor, N-butyldeoxynojirimycin was not successful in ameliorating neurological symptoms. However, it is relatively weak inhibitor of GCS with more potent inhibitory off-target effects (Schiffmann et al., 2008). However, more specific and potent GCS inhibitors are showing early promise (Schiffmann, 2020; Wilson et al., 2020).

## Acknowledgements

We thank Dr Stefan Karlsson for generously sharing lnl/lnl, nGD mice. We also thank Dr. Sreeganga Chandra for suggestions on mice behavioral tests.

## Material Methods

### Patients

Stored serum samples of Gaucher disease patients were analyzed. All patients had diagnosis of Gaucher disease based on <10% of normal leucocyte acid-β glucosidase activity. The study was approved by the Human Investigations Committee of Yale School of Medicine. 51 patients had type 1 Gaucher disease with at least one N370S (pArg409Ser) mutation in *GBA1* gene, mean age 63.9 years (range 40 to 93 yrs.). Patients were stratified into young and older patients with mean age 11.25 yrs. vs 55.8 yrs. Five patients had Gaucher disease type 3 (homozygous for L444P mutation, pLeu483Leu). Mean age of 28 healthy controls was 38 yrs.

### Mice

Mice were housed in the animal facility of Yale university in New Haven. All animal experiments were conducted in compliance with institutional regulations under authorized protocol (2016-10872) approved by the Institutional Animal Care and Use Committee. Gba^lnl/lnl^ mice or K14^lnl/lnl^ were generated as described previously described (Enquist et al., 2007). We referred Gba^lnl/lnl^ as nGD mice throughout the text. We used K14 Cre, Cx3cr1 Cre and Nestin Cre obtained from Jackson labs. For breeding purpose, we used Gba ^lnl/wt^ mice (gift from Sanofi Genzyme) that were then crossed with K14 ^Cre/Cre^ (Jackson labs) to obtain Gba ^lnl/wt^ K14 ^cre/cre^. These mice were used as parents to obtain nGD, Gba^lnl/wt^ and Gba^wt/wt^ pups. nGD Cx3cr1^Cre/+^ and nGD Nes^Cre/+^ were obtained by breeding these parents on to Cx3cr1 Cre and Nestin Cre. Gba^loxp/loxp^ Cx3cr1 ^cre^ mice were generated by crossing Gba^loxp/loxp^ (Mistry et al., 2010) with Cx3cr1 Cre mice obtained from Jackson labs. Mice of both sexes were used for the study

#### Brain tissue harvesting and cell preparation

The complete brain was cut in to small pieces and incubated with digestion buffer (RPMI supplemented with 2% FBS, 2mM HEPES, 0.4mg/ml Collagenase D and 2mg/ml DNase) for 30 min at 37°C under shaking. To stop enzymatic digestion EDTA 5mM was used and the sample was homogenized with a syringe. This was then followed by gradient centrifugation and cell separation was achieved via Percoll gradients (GE Healthcare Life Sciences) of various densities (Lee and Tansey, 2013). Myelin was then removed by vacuum suction and cells were isolated from the interphase. The interphase was diluted with 10ml of HBSS and centrifuged at 500g for 7 min to obtain cell pellet that was used for cellular analysis.

#### Flow cytometry

Flurochrome labelled antibodies used for the flow cytometry are listed in the key resource table. In brief surface staining was performed ex vivo using antibodies to CD11b, CD45, Gr-1, CD3, CD8, CD4, CD103, CD69 and NK1.1 (BD Biosciences); ACSA-2 antibody (Miltenyi Biotech). For staining of the intracellular antigens GZM-A and Pro IL-1β (BD Biosciences) cells were stimulated with cell stimulation cocktail (ebioscience) for 4h. After surface staining with antibodies, cells were fixed and permeabilized using BD cytofix and Cytoperm and antibodies recognizing GZM-A and Pro IL-1β.

### Sorting of cells by flow cytometry

Isolated total brain cells were resuspended at 1 million cells/ml of PBS and Live/Dead Fixable Violet Dead Cell Stain (ThermoFisher) for 10 min to exclude nonviable cells. Cells were washed once in excess PBS and then, cells were suspended in ice-cold FACS buffer (10% FBS in PBS with 1% HEPES +0.5% EDTA) and stained with anti-mouse CD45 and anti-mouse CD11b antibody (BD Pharmigen) for 30 min at 4 °C. All cells were washed twice with FACS buffer and sorted into polypropylene tubes with 500 μl of ice-cold FACS buffer. All samples were acquired on the BD FACS Aria.

#### Single-cell RNA sequencing (scRNA) seq

All cells were prepared through 10X Genomics V3 3^’^ Gene Expression kit and sequenced on NovaSeq flow cells to achieve high read depth. We used the pre-processed digital gene expression matrices as obtained from cell ranger (as run by the YCGA sequencing core facility). These matrices were processed using Seurat (Satija et al., 2015). The clusters were determined using Louvain algorithm for community detection (Vincent D Blondel, 2008). Differential gene expression between the clusters was carried out using the MAST method (Finak et al., 2015). The visualizations and analysis were carried out in R (R Core Team). The data is visualized as tSNE (Hinton, 2002). From 14day pups we sequenced total of 4895 CD45+ cells. For GZ161 treatment experiments we sequenced total of 40763 CD45+ cells that consists of 4 groups of mice with 3 mice/ each group.

#### Subsetting of microglia cells

Based on the clustering of CD45^+^ cells clusters 1, 16 and 26 were kept for final analysis. The clusters were selected based on the microglial gene expression. The clusters comprised 4230 cells. Scoring used z-scores of homeostatic microglia genes, DAM genes from (Wang et al., 2020).

### Single-nuclear RNA sequencing (snRNA seq)

#### Nuclei Isolation

Brain Tissue samples were stored at −80°C. For tissue lysis and washing of nuclei, sample sections were added to 1 mL lysis buffer (Nuclei PURE lysis buffer, Sigma) and thawed on ice. Samples were then Dounce homogenized with PestleAx20 and PestleBx20 before transfer to a new tube, with the addition of additional lysis buffer. Following incubation on ice for 15 minutes, samples were then filtered using a 30 μM MACS strainer (MACS strainer, Fisher Scientific), centrifuged at 500xg for 5 minutes at 4°C using a swinging bucket rotor (Sorvall Legend RT, Thermo Fisher), and then pellets were washed with an additional 1 mL cold lysis buffer and incubated on ice for an additional 5 minutes. Lysates were combined with 1.8ml of a 1.8M sucrose solution (Nuclei PURE Sucrose Buffer, Sigma) containing 1mM DTT and 0.2U RNAse inhibitor and mixed by inversion. Samples were then layered on top of a 1.8M sucrose layer to form a gradient and centrifuged at 30,000xg for 45min at 4°C using a swinging bucket rotor. Supernatant was removed, and nuclei pellets were resuspended in 1ml of a 1XPBS wash buffer containing 1% BSA and 0.2U RNAse inhibitor. Samples were centrifuged again at 500xg for 5 minutes at 4°C and supernatant removed. Nuclei were resuspended in 0.5ml wash buffer and counted using Countess (Life Technologies) prior to 10xGenomics protocol. For samples that were enriched using flow cytometry, nuclei were stained with 20 μg/ml DAPI for 30 minutes on ice with occasional mixing. Nuclei were centrifuged at 500xg for 5minutes at 4C. Supernatant was removed, and pellet resuspended in 800 μl of a 1XPBS wash buffer containing 1% BSA, 1mM EDTA and 0.2U RNAse inhibitor. Nuclei were sorted using the BD Influx by first gating on forward/side scatter then on DAPI positive nuclei. Collected nuclei were centrifuged then resuspended in ∼200ul 1XPBS wash buffer containing 1% BSA and 0.2U RNAse inhibitor and counted using Countess (Life Technologies) prior to 10xGenomics protocol.

#### Library preparation and NovaSeq Sequencing

Libraries were prepared according to 10xGenomics protocol using the Chromium Next GEM Single Cell 3’ Reagents Kit V3.1 (Dual Index) for encapsulation, mRNA capture, cDNA synthesis/amplification and library construction. Final libraries were quantified using the DNA High Sensitivity Kit (Agilent Bioanalyzer 2100) and Qubit 2.0 (Life Technologies). Libraries were diluted to 1.5nM then pooled prior to sequencing on the Illumina NovaSeq6000. UMI count matrices generated by Cellranger V3.0.2.

#### Data Preprocessing

Summary information for final UMI count matrices for nuclei by individual sample and sequencing data are presented in (Sup Fig 5A and C). Count matrices together with nucleus barcodes and gene labels were loaded with R version 3.6.1/RStudio for sample integration and unsupervised clustering using Seurat Package version 3.1. For Quality Control (QC), nuclei were filtered following standard protocols based on examination of violin plots. Cutoffs were used, filtered matrices were then individually log-normalized by sample according to standard Seurat workflows. After quality filtering, total nuclei were included in the final data analysis.

#### Broad Cell Types Annotation

Sample integration was performed in Seurat using the FindIntegrationAnchors and Integrate Data functions for variable features. Following integration and scaling according to Seurat package workflows, a range of clustering resolution values were trialed prior to broad cell type annotation; UMAP, TSNE plots for broad types annotation included. Cluster-level expression of major cell type markers was examined and used to annotate cells contained within each cluster. Clustering resolution achieved separation and consistency of cell type marker expression.

#### Differential Gene Expression and Pathway Enrichment Analysis

Genes differentially expressed between conditions within clusters may reveal pathway alterations specific to the clustered cell type. To profile cluster-level pathway enrichment patterns, cluster-level differentially expressed genes were identified using the Seurat Find Markers function with the MAST package. Markers used for downstream functional analysis were those with adjusted p-value < 0.05. Functional ontology analysis by cluster was performed for each cluster marker set using IPA (enrichment adjusted p-value < 0.05).

#### Total RNA seq

mRNA sequence data were uploaded to a high-performance computing system by PartekFlow software (v7.0, Partek, St. Louis, MO), adapter trimmed and remapped to mouse genome, mm10 using STAR v2.5.3a aligner with default setting (Phred:20) for read mapping. Statistical analysis was carried out using false discovery rate (FDR) correction through the Benjamin-Hochberg method.

#### BrdU retention assay

The assay was performed as described earlier (Takizawa et al., 2011), briefly mice were i.p injected with 180 μg BrdU (sigma) and were fed water containing 800 μg/ml BrdU and 4% glucose for 12days. Mice were then sacrificed organs (brain and spleen) were removed and cells were isolated as described earlier. BrdU staining was performed using BrdU labelling kit (BD). Cells from normal water fed mice were used as staining controls.

#### Mouse serum NF-L assay

Quanterix Nf-L assays were performed in triplicate according to the manufacturer’s protocols using the Nf-Light^TM^ Advantage kit on a single molecule array (SIMOA^TM^) HD-X instrument (Quanterix, Lexington, MA).

#### Lipidomics

Separation of glucosylceramides and galactosyl ceramides was performed by SFC-MS/MS analyses at the MUSC Lipidomics Shared Resource. The equipment consisted of a Waters UPC 2 system coupled to a Thermo Scientific Quantum Access Max triple quadrupole mass spectrometer equipped with an ESI probe operating in the multiple reaction monitoring MRM positive ion mode tuned and optimized for the Waters UPC 2 system.

#### Brain tissue slide preparation and MALDI-MS imaging

Brain tissues were dissected, divided sagittally into halves and immediately placed in Cryomolds (Tissue-Tek) containing 10% gelatin-water (porcine skin, Sigma-Aldrich, #G1890), placed at 37C on the heating block and transferred the cryo-mould in to dry-ice bath. Cryo-sectioning was performed at a chamber temperature of −20 °C with 12 µm thickness. Sections were thaw-mounted onto the ITO coated side with barcoding of MALDI IntelliSlides (Bruker, cat#1868957). Slides were stored in −80 °C until time for imaging. An optical image for tissue sections was obtained using TissueScout scanner (Bruker). Once the optical image was obtained, it was transferred to the HTX Sublimator (HTX Technologies, Chapel Hill, NC) for matrix deposition. DHB (2,5-dibydroxybenzoic acid) was applied on the tissue sections by sublimation, which was performed according to the HTX Sublimator setting (2ml of 40mg/mL DHB in acetone transfer solvent, 60 °C preheat temperature, 160 °C final temperature, 200 seconds). MALDI-MS imaging was performed in positive ion mode over a mass range of m/z 300-1300 on a Bruker timsTOF fleX mass spectrometer equipped with a 10kHz SmartBeam 3D Nd:YAG (355mm) laser. Imaging was performed using a laser raster size of 20 µm custom setting, 20 µm scan range, with trapped ion mobility mode On (tims’ON’) for cross-sectional collision (CCS) values with the following parameters (1/K_o_ start at 0.90 V.s/cm^2^ and end 1.70 V.s/cm^2^, ramp time at 150 ms, accumulation time 40.0 ms, duty cycle 26.67%, ramp rate 6.50 Hz). Spectra were accumulated from 400 laser shots with the laser power percentage adjusted using the highest peak (m/z 760 positive ion mode) intensity between 10^4^ and 10^5^, and the method was saved for all subsequent acquisitions. For each image acquisition, the instruments were calibrated using ESI-L Low concentration tuning mix (Agilent Technologies, cat #G1969-85000). Data were analyzed using SCiLs Lab MALDI imaging software package, version 2021b, (Bruker) with data normalized to the TIC. Targeted analysis for Hexceramides, sphingomyelins, and phosphatidylcholines were analyzed for differential changes between animals.

### Lipid extraction of mouse serum

To quantify GlcSph, 7µL of mouse serum was aliquoted into a labeled 1.5mL Eppendorf tube followed by 100 µL of internal standard solution (12ng/mL dimethyl-psychosine, 80% methanol, 20% acetonitrile with 10mM ammonium acetate and 1% formic acid). The samples were vortexed for 5mim and sonicated for 10min. The tubes were centrifuged at 13,000g for 5min at room temp. The supernatant from each tube was transferred into a pre-labeled total recovery MS vial. Calibration curves for GlcSph was prepared in a pooled control serum, and concentrations of lysoGL1 are from 0.1–1000 ng/mL.

### Mass spectrometry analysis of GlcSph

Mouse serum was first tested on a Hilic-method to separate psychosine (steroisomer of GlcSph) and GlcSph and confirm that psychosine does not interfere with the quantitation of mouse serum glucosylsphingosine levels (Wei-Lien Chuang etal). The supernatant was injected (5 µL) into an LC/MS/MS system comprised of an Acquity UPLC (Waters, Milford, MA) and Sciex Triple Quad 5000 mass spectrometer (Sciex, Toronto, Canada). The chromatographic separation was achieved with an Waters Acquity BEH C18 column (2.1 ∗ 150 mm, 1.7 µm) using mobile phases: (A) Water with 0.1% formic acid, (B) 85:15 MeOH:ACN with 0.1% formic acid. The column is maintained at 60 C. GlcSph is eluted with the following gradient: from 50% B to 99% B over 2 min, then the mobile phase composition is hold constant for 1 min followed by a rapid return (0.1 min) to 50% B maintained for 0.5 min. All experiments were carried out at a flow rate of 0.5 mL/min. Data is analyzed in Analyst (AB Sciex, Toronto, Canada).

### Behavioral Studies

Balance beam test: Evaluating fine motor coordination using balance beam (1mt) with 12mm width resting 50 cm above the top of the pole. The time to cross each beam was recorded and compared between mouse strains.

### Open Field

Mouse locomotor behavior was assessed using the open field (Crusio, 2001). Mice were individually placed on the 28×28 cm plate surrounded by plastic walls in a well-lit room and allowed to freely explore for 10 min. Spatial statistic, total distance travel, area measure, and time spent in the center of the square were quantified. The movements were recorded by a video tracking system and stored on a computer.

### BODIPY staining

Isolated total mice brain cells from nGD and age matched control mice were prepared (as described in section “Brain tissue harvesting and cell preparation”) were stained with CD45, CD11b and ACSA-2 were incubated in PBS with BODIPY 493/503 (1:1,000 from a 1 mg ml stock solution in DMSO; Thermo Fisher) for 10 min at RT, washed twice in PBS, and BODIPY intensity was analyzed on LSRII instrument (BD Biosciences).

### ROS assay

To assess ROS generation in astrocytes and microglia from nGD mice, isolated total mice brain cells were stained with CD45, CD11b and ACSA-2 followed by incubation in FACS buffer with CellROX Deep Red (1:500; Invitrogen) for 30 min at 37 °C, washed twice in FACS buffer, and CellROX Deep Red Intensity was analyzed on LSRII instrument (BD Biosciences).

### ApoE measurement

Apoe levels in sera of GD patient were measured by ELISA using Abnova CAT # KA1031 following manufacturer’s instructions. A dilution of 1:400 sera was used.

### Quantification and Statistical analysis

Data were routinely presented as the mean ± SEM. Statistical significance was determined using t test using Bonferroni-Dunn correction for multiple comparisons. Differences between groups were analyzed using Student’s t test using GraphPad Prism 8.0. For the Kaplan-Meier analysis of survival, the log-rank (Mantel Cox) test was performed.

### Study approval

All animal experiments were conducted in compliance with institutional regulations under authorized protocol (2016-10872) approved by the Institutional Animal Care and Use Committee. The study was approved by the Human Investigations Committee of Yale School of Medicine and written informed consent was received prior to participation.

| Reagents | Provide supplier name | Catalogue number | Clone |
| --- | --- | --- | --- |
| CD11b | BD Biosciences | 583553 | M1/70 |
| CD11b | BD Biosciences | 552850 | M1/70 |
| ACSA-2 | Miltenyi Biotech | 130-123-284 | IH3-18A3 |
| CD45 | Biolegend | 103133 | 30-F11 |
| NK1.1 | BD Biosciences | 562864 | PK136 |
| CCR2 | R and D | FAB5538F |  |
| CD3 | Biolegend | 100241 | 17A2 |
| NK1.1 | Biolegend | 108753 | PK136 |
| 1-A/1-E | Biolegend | 107639 | M%/114.152 |
| CD4 | BD Biosciences | 563790 | GK1.5 |
| CD4 | BD Biosciences | 558107 | RM4-5 |
| Gr-1 | Biolegend | 108440 | RB6-8c5 |
| Ly-6C | Invitrogen | 47-5932-82 | HK1.4 |
| CD64 | Biolegend | 139306 | X54-5 |
| B220 | eBioscience | 47-0452-80 | RA3-6B2 |
| CD8a | eBioscience | 56-0081-82 | 53-6.7 |
| Brdu Flow kit | BD Pharmingen | 51-9000019AK |  |
| 5-Bromo-2'-deoxyuridine | Cayam Chemical Company | 15580 |  |
| Cell Rox Deep Red | Invitrogen | C10422 |  |
| IL-1Beta | Invitrogen | 17-7114-80 | NTTEN3 |

### Resource availability

#### Lead contact

Further information and requests for reagents should be directed to and will be fulfilled by the lead contact, Pramod Mistry.

#### Materials availability

All materials in this study are available upon request.

#### Data and code availability

All data produced in this study are included in this published article and the supplemental information. Any additional information required to reanalyze the data reported in this paper is available from the lead contact upon request.

## Author contributions

C.S.B, S.N. conceived the project, performed, designed experiments and analyzed data; G.B, J.R involved in mouse generation and maintenance; E.P, S. M performed bioinformatics analysis; J.S performed the snRNA-seq analysis; V.D.J performed mice behavioral studies; T.N, performed MALDI-MS imaging; L.G, M.K, H.W performed Quanterix assays and lipidomics in mice and human serum. M.D, Z.B and K.L provided helpful insights. P.K.M. conceived the project, supervised the study and analyzed data; C.S.B, S.N and P.K.M wrote the manuscript with input from all authors.

---

**Figure S1.**
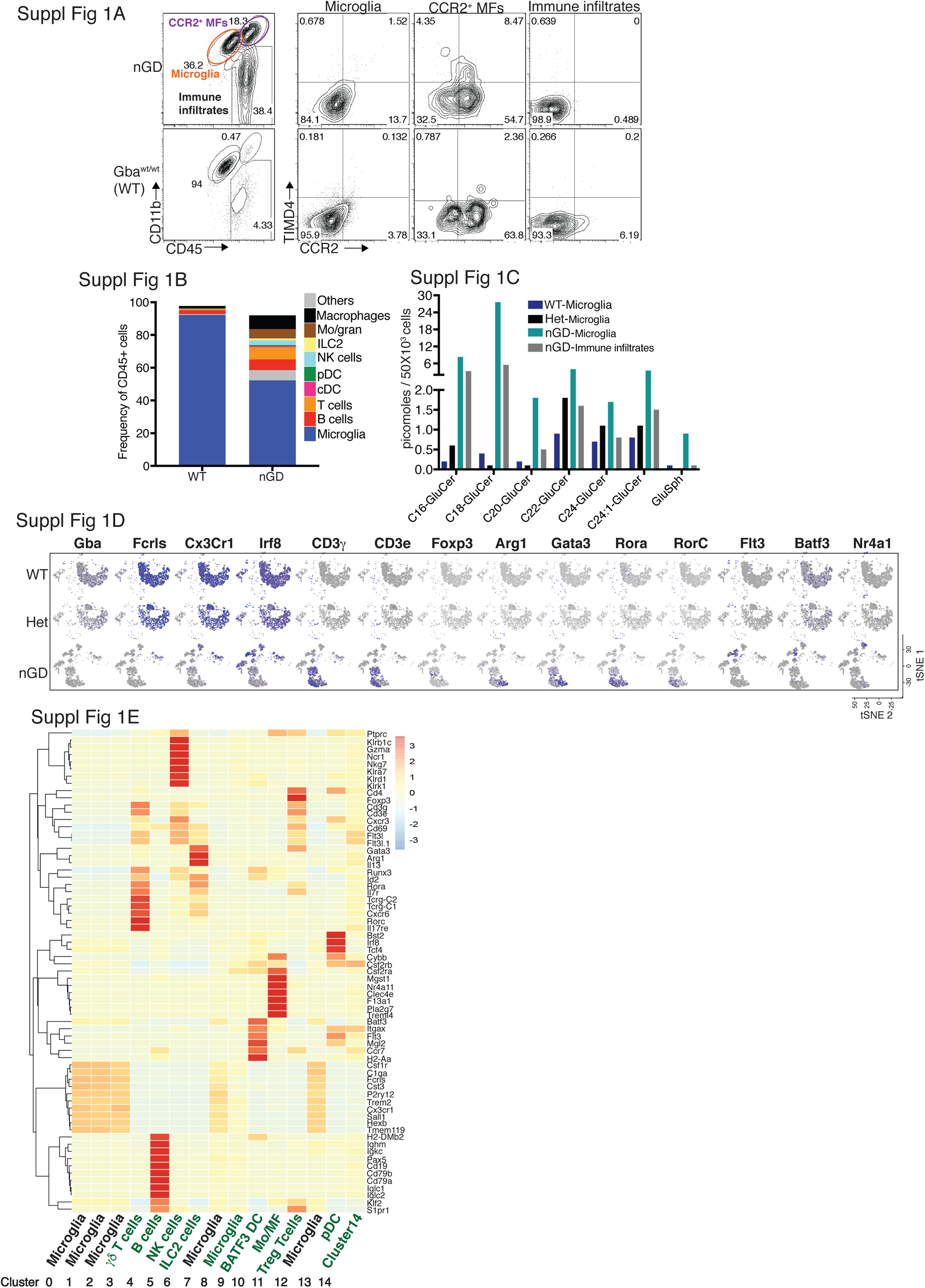
Loss of Gba induces microglial activation and immune cell infiltration in neuronopathic GD (Gba lnl/lnl) brain. A. Representative FACS plot showing CCR2+ MFs, microglia and immune infiltrates in the whole brain of nGD versus control mice brain.B. Bar graph shows the frequency of different immune cells (denoted on right) in the immune infiltrates obtained from neuronopathic GD (nGD;Gbalnl/lnl) and control mice brain respectively. C. Bar graph shows quantitative analysis of total GluCer species and GlcSph levels by LC-ESI-MS/ MS in flow sorted microglia cells obtained from Gbawt/wt, Gbalnl/wt and nGD brains and on immune infiltrates isolated from nGD brain. D. t-SNE plot demonstrating annotations of different clusters identified using expression of signature genes. E. Hierarchical heatmap representing the differentially expressed genes in each identified cluster.

**Figure S2.**
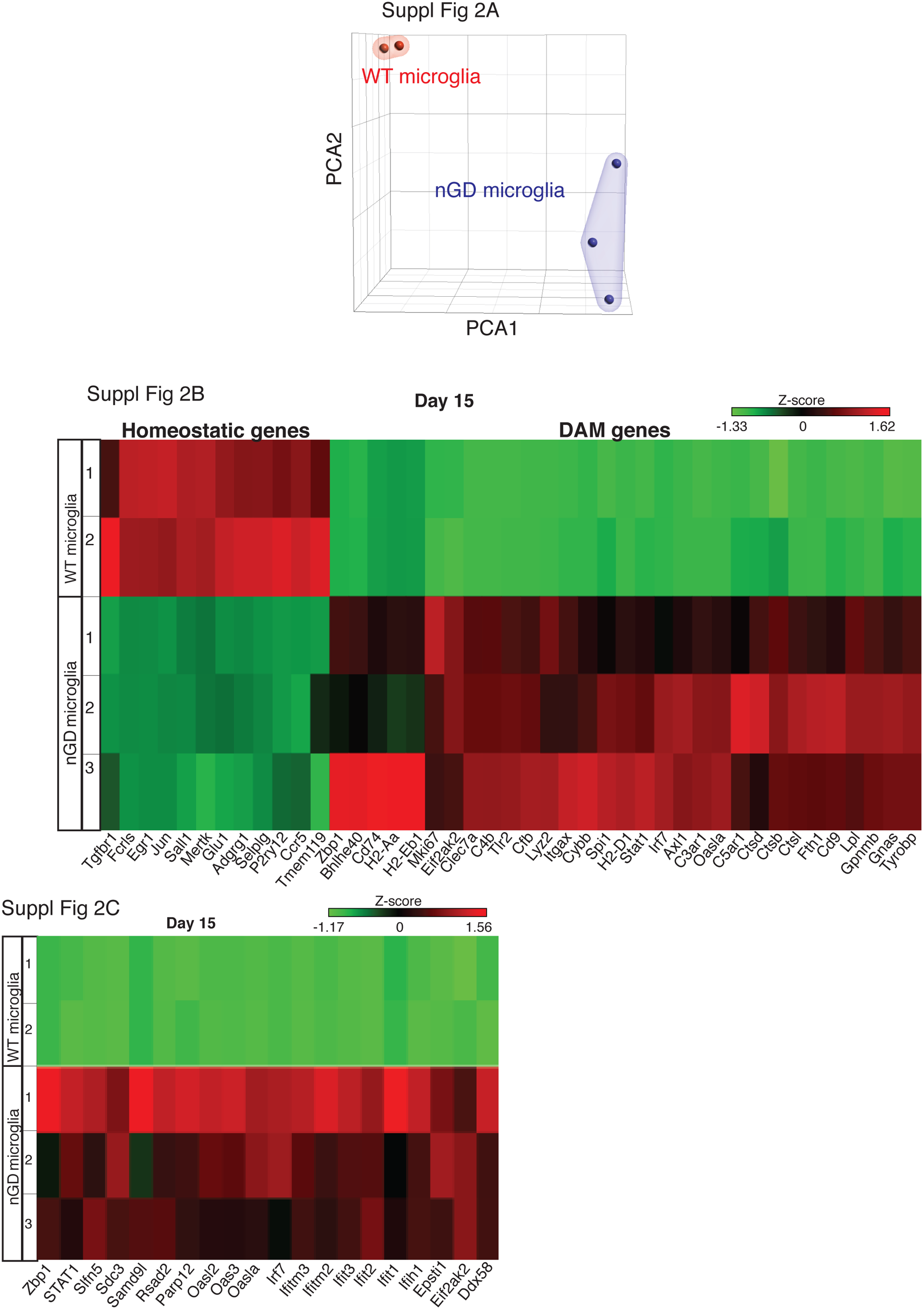
Loss of Gba disrupts microglial homeostasis and induces damage associated microglia (DAM) phenotype A. Principal component analysis of gene expression profiles of microglia isolated from nGD brain and their controls. Each point represents a single mouse. B. A heat map showing the differential expression of homeostatic and Disease associated microglial (DAM) genes in isolated-microglia from nGD and control mice brain respectively. C. A heat map of interferon signature genes (ISG) genes in isolated-microglia from nGD and control mice brain respectively. Colors indicate upregulated (red) and downregulated genes (green).

**Figure S3.**
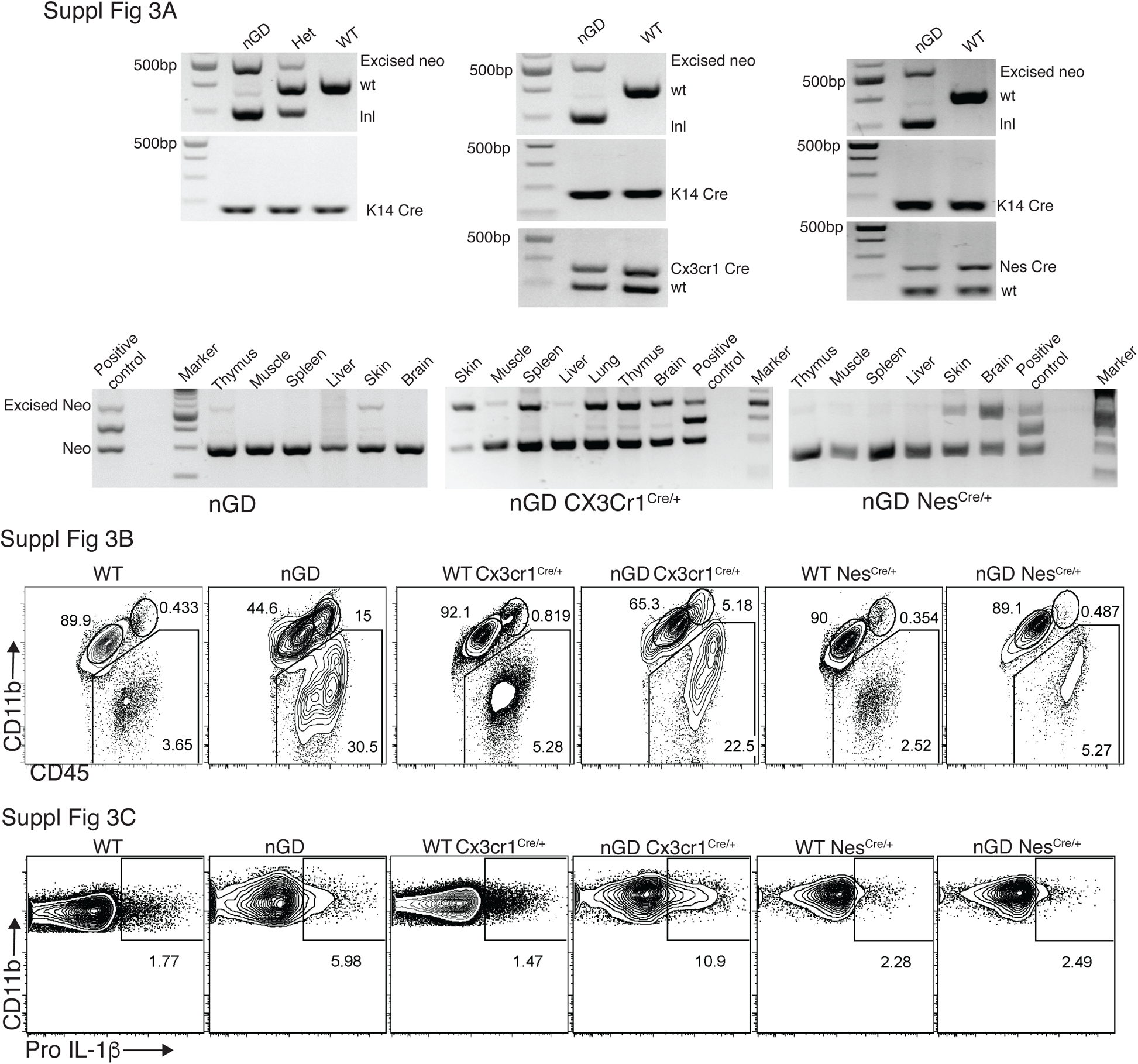
Restoring Gba function in microglia and neurons of enhances nGD mice survival. A. Splicing of mRNA causes GBA1 deficiency in nGD mice. K14-mediated expression of Cre enabled removal of the lnl (loxp-neo-loxp) cassette in the skin tissue of the nGD mice (left and bottom panel), Cx3cr1 Cre enabled removal of the lnl cassette in different tissues (middle and bottom panel) and Nestin Cre enabled removal of the lnl cassette in brain (right and bottom panel). B. Representative FACS plot showing CCR2+ MFs, microglia and immune infiltrates in the whole brain of nGD, nGD Cx3cr1Cre/+ and nGD NesCre/+ mice versus control mice brain. C. Representative intracellular staining for Pro-IL-1ß expression in microglia isolated from nGD, nGD Cx3cr1Cre/+ and nGD NesCre/+ and control mice analyzed by FACS.

**Figure S4.**
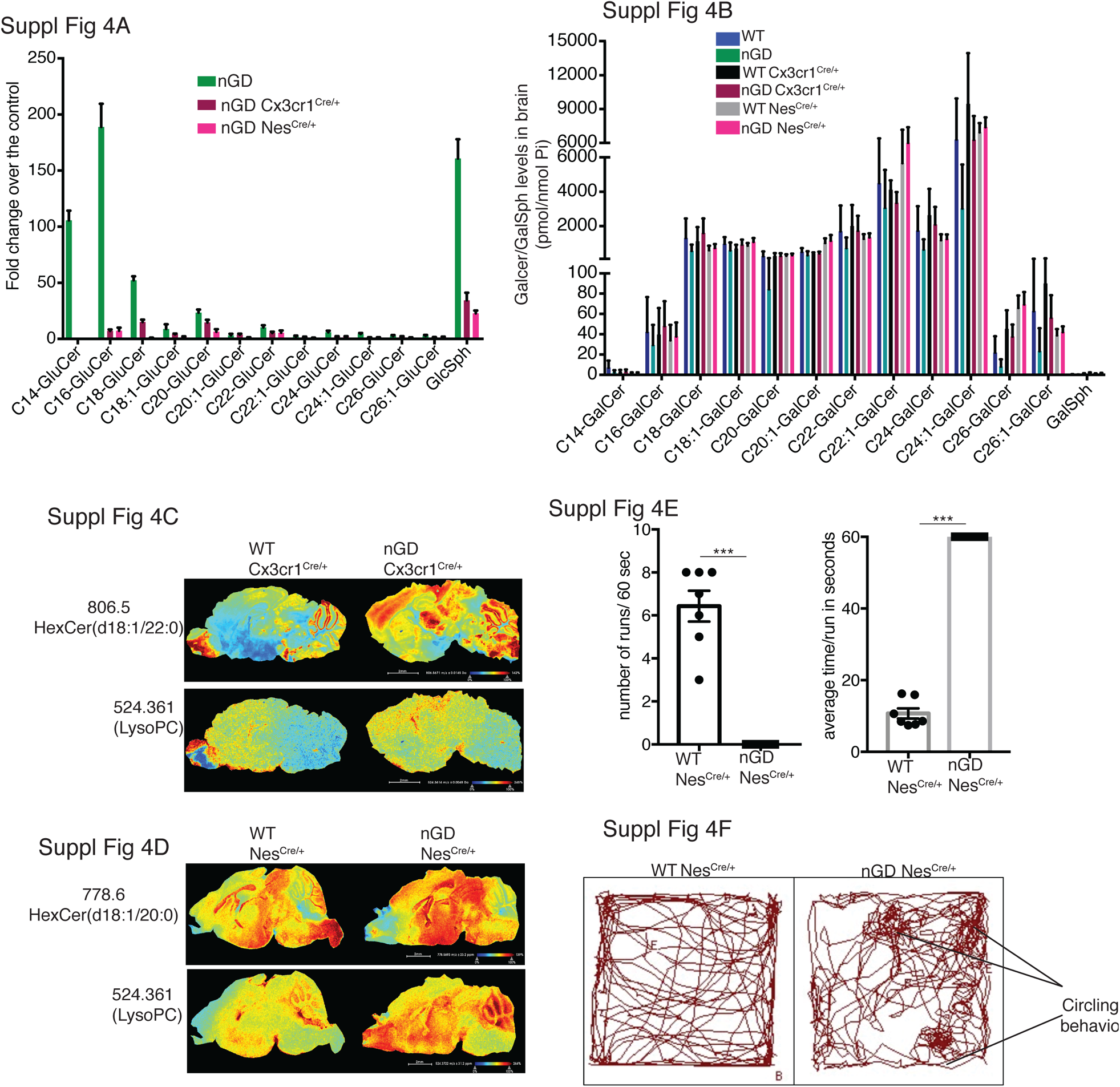
*Gba* deficiency in neurons aid in microglia activation and immune cell infiltration. A. Fold change increase in different Glucosylceramide (GluCer) and glucosylsphingosine (GlcSph) species in nGD, nGD Cx3cr1^Cre/+^ and nGD Nes^Cre/+^ mice over respective control mice brain. **B.** Quantitative analysis of total GalCer species and GalSph levels by LC-ESI-MS/ MS in nGD, nGD Cx3cr1^Cre/+^ mice and nGD Nes^Cre/+^ mice brain compared with the control mice (n=4-8 mice/group). **C.** Signal intensities of HexCer species (d18:1/20:0) and LysoPC identified by MALDI across nGD Cx3cr1^Cre/+^ and control littermates **D.** and nGD Nes^Cre/+^ mice and control mice. The color bars in MALDi images show signal intensity: blue to red indicates low to high levels. The data is representative of three independent experiments. **E.** nGD Nes^Cre/+^ mice showed motor deficits in the balance beam test as compared to control mice. **F.** Representative track plots obtained from open-field studies show circling behavior in nGD Nes^Cre/+^ mice.

**Figure S5.**
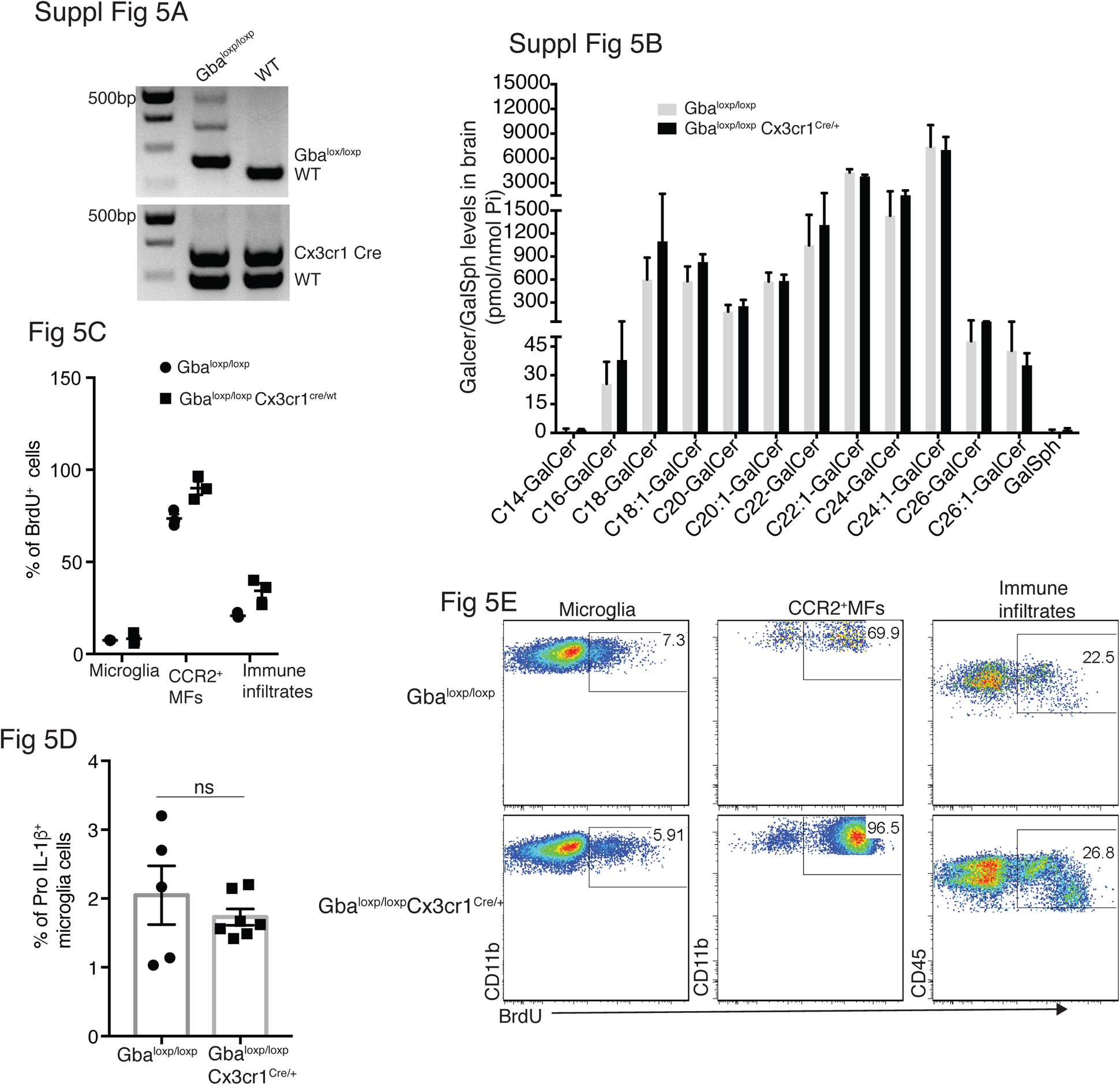
Changes in microglia subsets coupled with neurodegeneration is seen in aged Gbaloxp/loxp Cx3cr1Cre/+ mice brain. A. Genotyping for Cx3cr1 cre and loxp alleles. B. Quantitative analysis of total GalCer species and GalSph levels by LC-ESI-MS/ MS in Gbaloxp/loxp Cx3cr1Cre/+ mice brain compared with the control mice (n=4-8 mice/group). C. Comparison of percentage of BrdU+ microglia, CCR2+ MFs and immune infiltrates between Gbaloxp/loxp Cx3cr1Cre/wt and control mice brain (n=3 mice/group) D. Representative histograms depicting BrdU incorporation in the microglia, CCR2+ MFs and immune infiltrates isolated from Gbaloxp/loxp Cx3cr1Cre/wt and control mice brain. E. Percentage of Pro-IL-1ß+ microglia cells in control and Gbaloxp/loxp Cx3cr1Cre/wt mice (n=5-6 mice/group).

**Figure S6.**
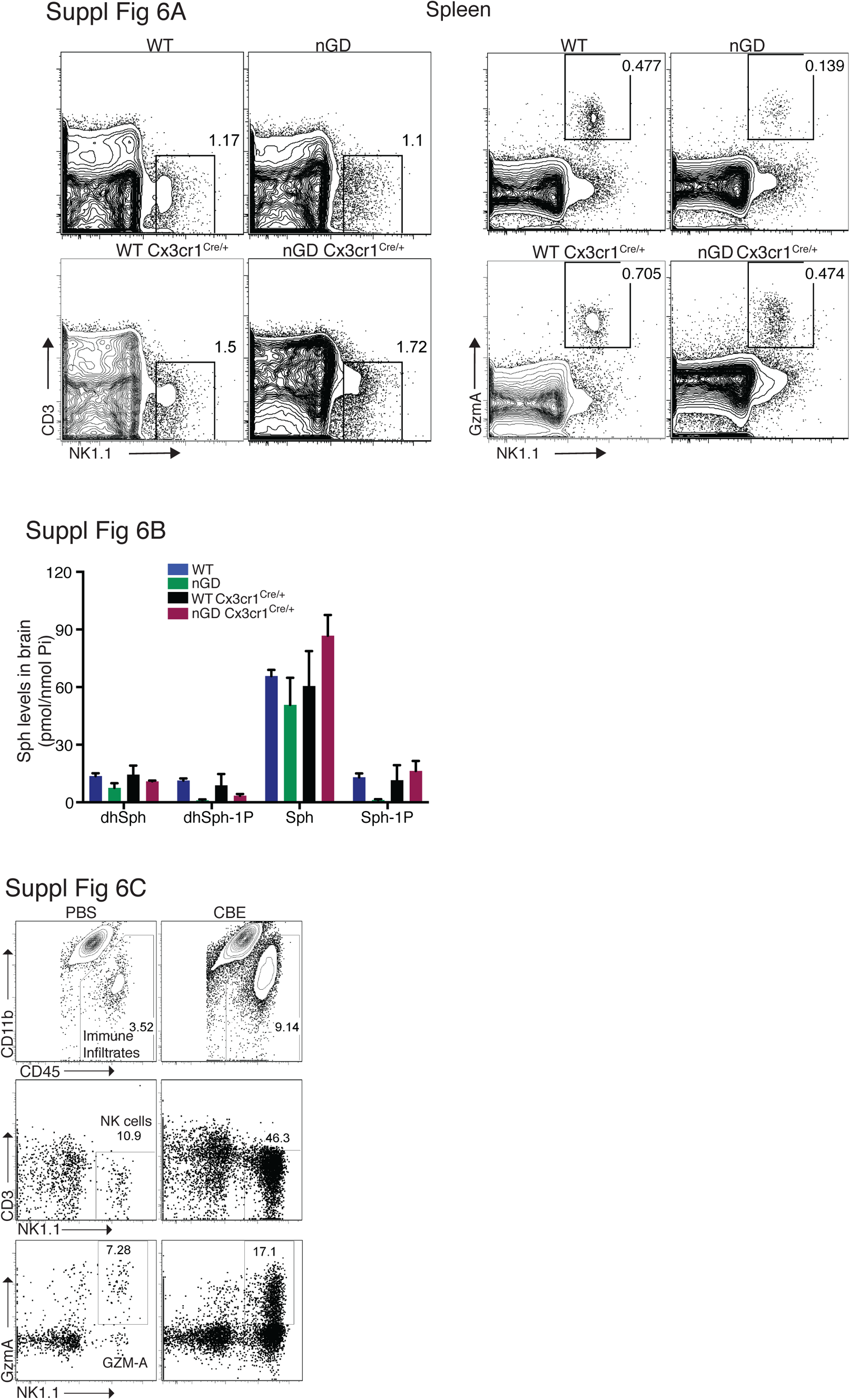
NK cell infiltration into the brain of nGD, nGD Cx3cr1Cre/+ mice. A. Representative FACS plot showing NK1.1+ NK cells and Gzm-A+ NK cells in the spleen of nGD, nGD Cx3cr1Cre/+ and control littermates. B. Lipidomic analysis by LC-MS/MS of Sphingosine-1-phosphate (S1P) and Sphingosine (Sph) content in brains from nGD, nGD Cx3cr1Cre/+ mice and control littermates. C. Representative FACS plot showing percentage of immune infiltrates (top panel), NK1.1+ NK cells (middle panel) and GzmA+ NK cells (bottom panel) in B6 treated with vehicle or CBE.

**Figure S7.**
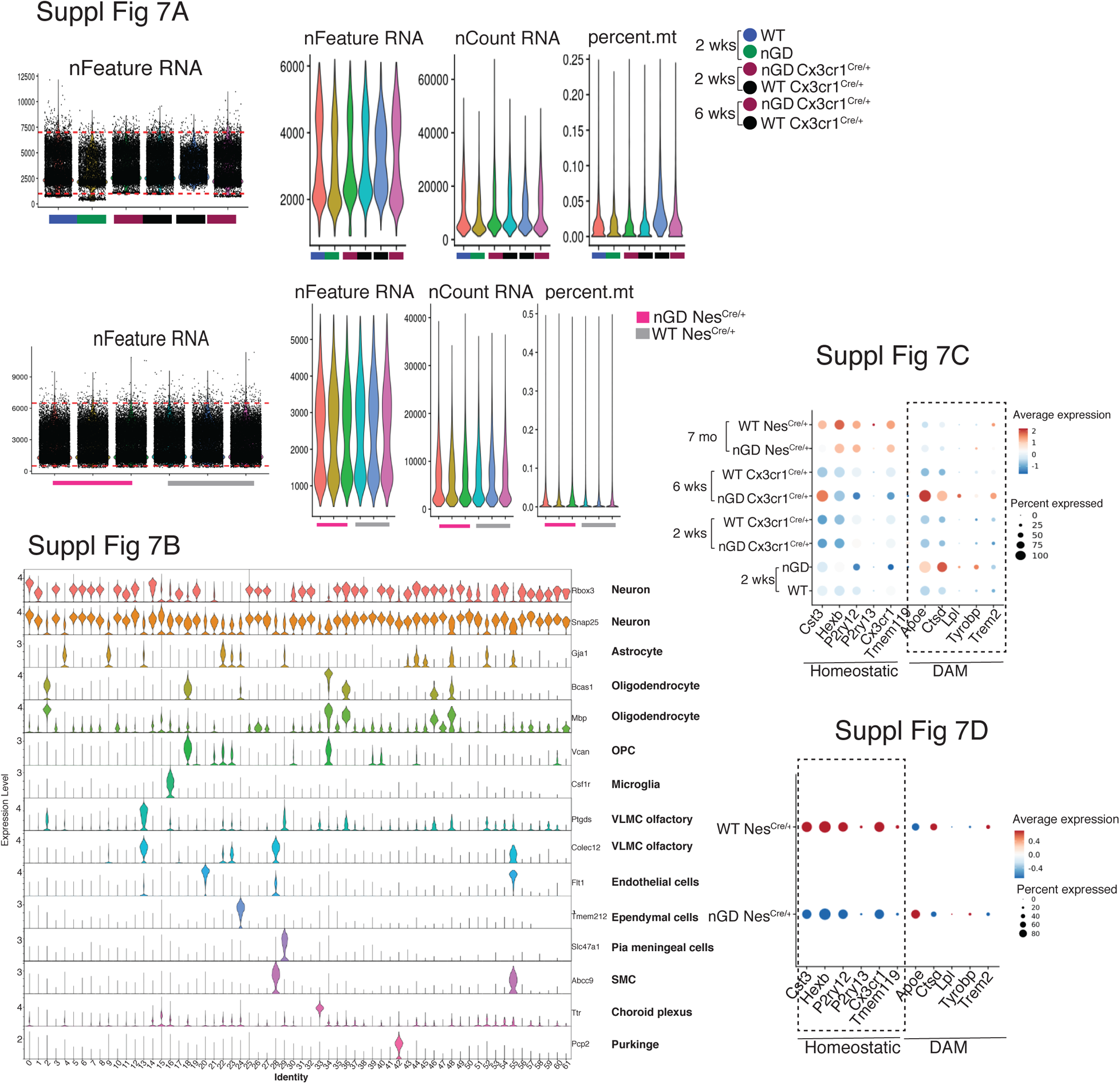
snRNA-seq analysis of the brain of nGD, nGD Cx3cr1Cre/+ mice. A. Analysis on snRNA-seq data of nGD (green), Gbawt/lwt (blue), nGD Cx3cr1Cre/+ 2 and 6 wks old (maroon), Gbawt/wt Cx3cr1Cre/+ 2 and 6 wks old (black), nGD NesCre/+ mice (pink) (n=3) and Gbawt/wt NesCre/+ (grey) (n=3). Violin plots showing the total number of detected genes (nFeature), reads counts (nCount), proportion of mitochondria (percent.mt) contamination per nucleus for each sample. B. Violin plots showing the cluster-specific expression of the canonical marker genes across all clusters. C. Dot plot showing the differential expression of homeostatic and Disease associated microglial (DAM) genes (dotted box) in microglia from nGD, Gbawt/lwt, nGD Cx3cr1Cre/+ 2 and 6 wks old, Gbawt/wt Cx3cr1Cre/+ 2 and 6 wks old, nGD NesCre/+ mice (n=3) and Gbawt/wt NesCre/+ (n=3) respectively. D. Dot plot showing the differential expression of homeostatic (dotted box) and Disease associated microglial (DAM) genes in microglia from nGD NesCre/+ mice (n=3) and Gbawt/wt NesCre/+ (n=3) respectively.

**Figure S8.**
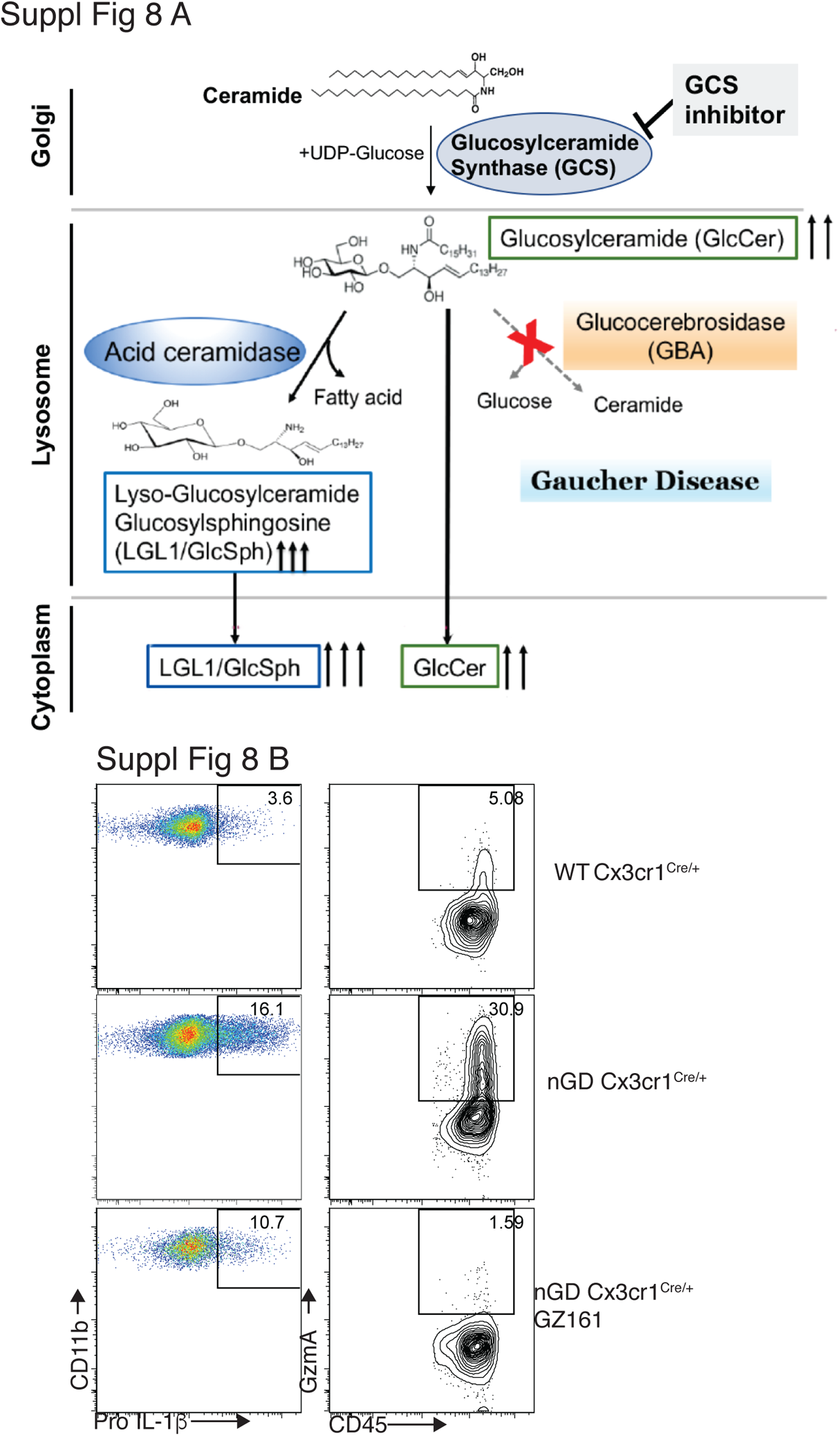
Immune regulatory effects of glucosyl ceramide synthetase (GCS) inhibitor GZ-161 on microglia and GzmA+ cells. A. Schematic illustration of mode of action of GCS inhibitor. B. Representative FACS plot showing percentage of Pro-IL-1ß+ microglia cells (left) and GzmA+ CD45+ (right) in wild type and nGD Cx3cr1Cre/+mice with and without treatment with GZ-161.

**Figure S9.**
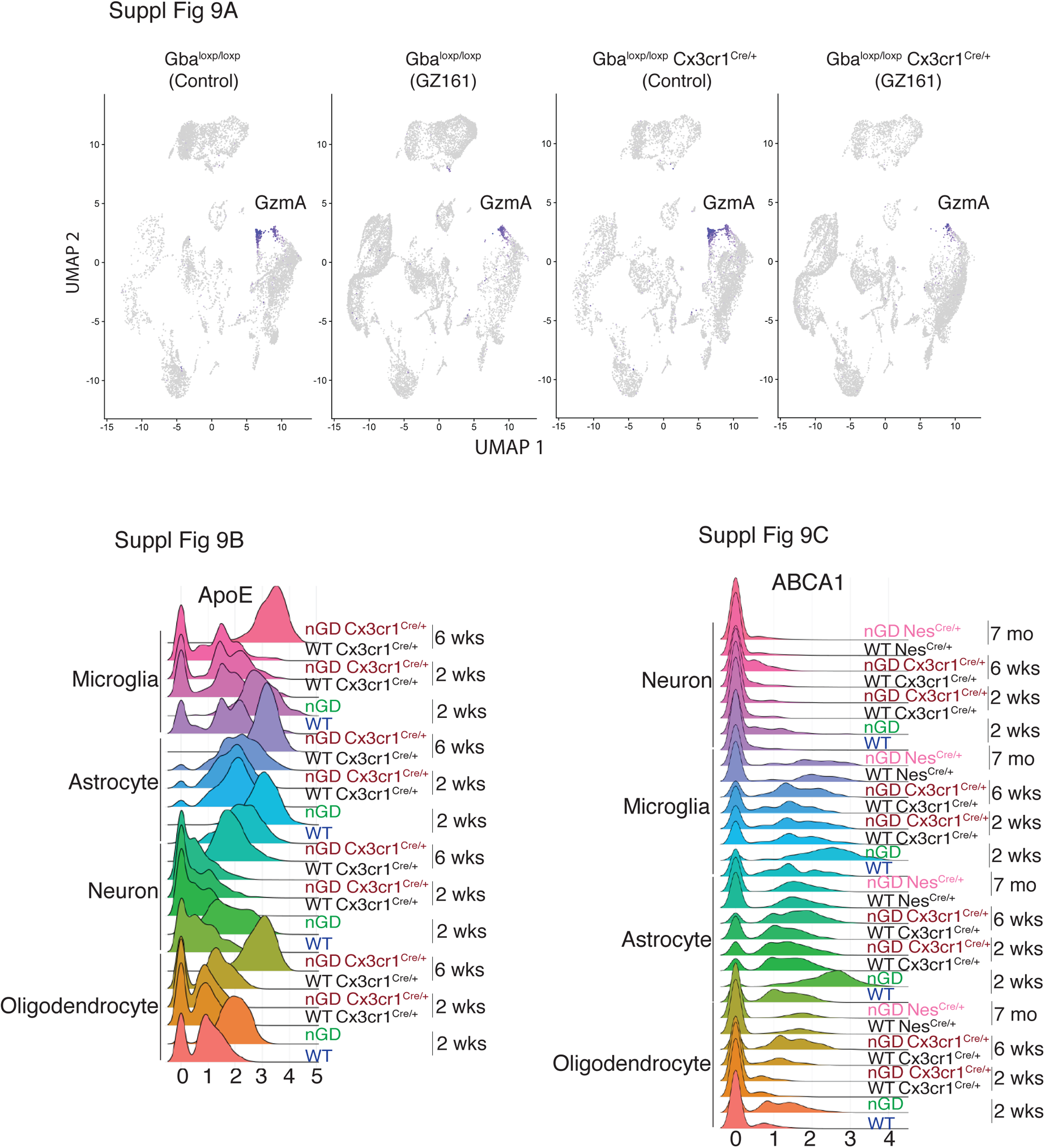
Long term treatment with GCS inhibitor, GZ-161, counteracts GluCer accumulation, improves microglial homeostasis, abrogates NK cell activation in Gba deficient microglia. A. UMAP showing GzmA expression in the clusters from control and Gbaloxp/loxp Cx3cr1Cre/wt mice treated with either vehicle or GZ161. B. Histogram showing differential expression of ApoE in microglia, astrocyte, neuron and oligodendrocyte cluster from nGD (2 wk. old), nGD Cx3cr1Cre/+ (2 and 6 wk. old respectively), and corresponding control mice. C. Histogram showing expression of Abca1 in oligodendrocyte, microglia, astrocyte and neuron from nGD (2 wk old), nGD Cx3cr1Cre/+ (2 and 6 wk old respectively), nGD NesCre/+ and corresponding control mice.

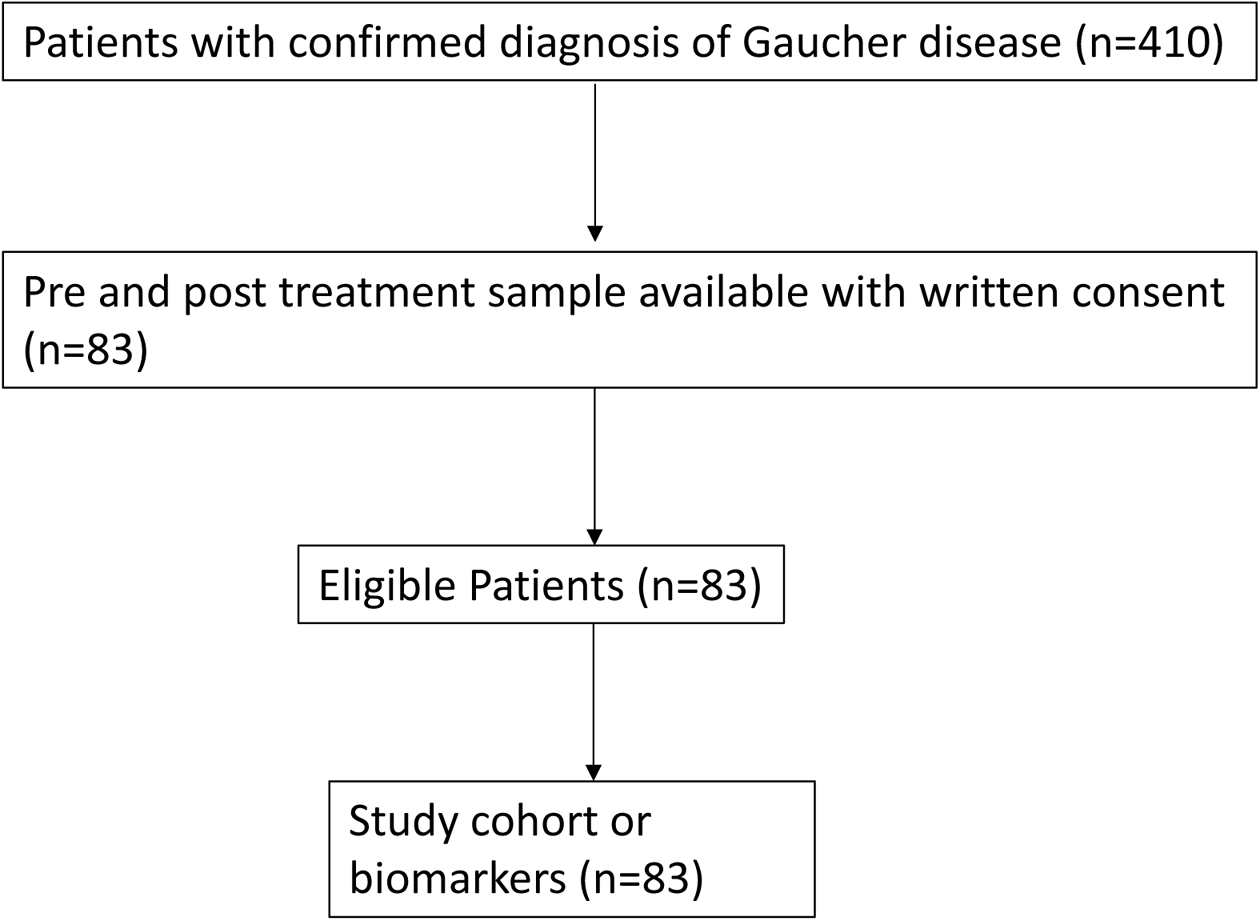

